# Generation of glucocorticoid resistant SARS-CoV-2 T-cells for adoptive cell therapy

**DOI:** 10.1101/2020.09.15.298547

**Authors:** Rafet Basar, Nadima Uprety, Emily Ensley, May Daher, Kimberly Klein, Fernando Martinez, Fleur Aung, Mayra Shanley, Bingqian Hu, Elif Gokdemir, Mayela Mendt, Francia Reyes Silva, Sunil Acharya, Tamara Laskowski, Luis Muniz-Feliciano, Pinaki Banerjee, Ye Li, Sufang Li, Luciana Melo Garcia, Paul Lin, Hila Shaim, Sean G. Yates, David Marin, Indreshpal Kaur, Sheetal Rao, Duncan Mak, Angelique Lin, Qi Miao, Jinzhuang Dou, Ken Chen, Richard Champlin, Elizabeth J. Shpall, Katayoun Rezvani

## Abstract

Adoptive cell therapy with viral-specific T cells has been successfully used to treat life-threatening viral infections, supporting the application of this approach against COVID-19. We expanded SARS-CoV-2 T-cells from the peripheral blood of COVID-19-recovered donors and non-exposed controls using different culture conditions. We observed that the choice of cytokines modulates the expansion, phenotype and hierarchy of antigenic recognition by SARS-CoV-2 T-cells. Culture with IL-2/4/7 but not other cytokine-driven conditions resulted in >1000 fold expansion in SARS-CoV-2 T-cells with a retained phenotype, function and hierarchy of antigenic recognition when compared to baseline (pre-expansion) samples. Expanded CTLs were directed against structural SARS-CoV-2 proteins, including the receptor-binding domain of Spike. SARS-CoV-2 T-cells could not be efficiently expanded from the peripheral blood of non-exposed controls. Since corticosteroids are used for the management of severe COVID-19, we developed an efficient strategy to inactivate the glucocorticoid receptor gene (*NR3C1*) in SARS-CoV-2 CTLs using CRISPR-Cas9 gene editing.

## INTRODUCTION

The emergence of severe acute respiratory syndrome coronavirus 2 (SARS-CoV-2) in 2019, marks the third and most devastating large-scale epidemic of coronavirus infection (known as COVID-19) in recent times. A number of potential treatment options against SARS-CoV-2 are under investigation, including the use of convalescent plasma, remdesivir, lopinavir/ritonavir, and interferon-beta(Beigel et al., 2020; Cao et al., 2020; Casadevall and Pirofski, 2020; Hung et al., 2020; NCT04315948, 2020). So far, none appear to be curative, making it critical to develop novel therapeutic strategies.

SARS-CoV-2 infection is characterized by profound T-lymphopenia associated with a dysregulated/excessive innate response, thought to be the underlying mechanism for acute respiratory distress syndrome (ARDS), the major cause of morbidity and mortality with this virus (Fathi and Rezaei, 2020; Mehta et al., 2020). Recent studies from patients with COVID-19 point to an important role for T cell adaptive immunity in protection and clearance of the virus (Braun et al., 2020), with T cell responses documented against the structural SARS-CoV-2 viral proteins spike (S), membrane (M) and nucleocapsid (N) (Grifoni et al., 2020; Ni et al., 2020; Sekine et al., 2020; Thieme et al., 2020). Indeed, in preclinical models of SARS-CoV-1 infection, adoptive transfer of virus-specific T cells (VSTs) was shown to be curative in infected mice, supporting the use of adoptive cell therapy (ACT) in coronavirus-related infection. ACT with allogeneic cytotoxic T-lymphocytes (CTLs) has been successfully used to treat other severe viral infections, such as cytomegalovirus (CMV), adenovirus, BK virus, Epstein-Barr virus and human herpes virus 6 in immunosuppressed patients, with responses ranging from 60-100% (Haque et al., 2007; Muftuoglu et al., 2018; O’Reilly et al., 2016; Tzannou et al., 2017). Thus, ACT may be an attractive approach for the management of COVID-19-related disease. However, many patients with severe COVID-19 receive corticosteroids which, due to their lymphocytotoxic effects, limit the efficacy of ACT. Here, we describe a novel approach for the generation of highly functional and steroid-resistant SARS-CoV-2 reactive T cells for the immunotherapy of patients with COVID-19.

## RESULTS

### Expansion of SARS-CoV-2 reactive T cells from COVID-19-recovered donors

Our group has previously reported the feasibility of generating VSTs from the peripheral blood (PB) of healthy donors for ACT (Muftuoglu et al., 2018). Here, we utilized this approach to derive and expand SARS-CoV-2 specific T-cells. Briefly, PBMCs from 10 CoV19-RD were cultured with 11 different peptide libraries (15mers overlapping by 11 amino acids) spanning the entire sequence of the SARS-CoV-2 antigens, including both the structural (S, M, N, E) and non-structural proteins (AP3A, Y14, NS6, NS7a, NS7B, NS8, ORF9B and ORF10) in the presence of either IL-2/4/7, IL-2/7/15, IL-2/4/21 or IL-2/7/21 for 14 days. At the end of the culture period, SARS-CoV-2 reactive T-cells were enumerated based on their ability to produce IFN-γ in response to *ex vivo* stimulation with the viral antigens. When cultured in the presence of IL-2/4/7 or IL-2/7/15, expansion was successful in 8/10 cases, with a median fold expansion of 719.14 (range 7.16 – 45572.50) and 1138.41 (range 15.97 – 27716.61), respectively. However, expansion using IL-2/4/21 or IL-2/7/21 was suboptimal, with a median fold expansion of only 0.71 (range 0.08-996.18) and 2.72 (range 0.85 – 415.98), respectively (**Figure 1A, Tables 1, S2 and S3**). IL-2/4/7 and IL-2/7/15 culture conditions supported expansion of both CD4+ and CD8+ SARS-CoV-2 specific T-cells with a predominance of CD4+ T-cells, while expansion with IL-2/4/21 and IL-2/7/21 failed to result in significant expansion of either SARS-CoV-2 CD4+ or CD8+ T cells (**Figure 1B**).

**Table 1.** Cytokine production of SARS-CoV-2 T cells from recovered donors expanded with different cytokine cocktails against M, N and S structural proteins.

|  |  | CD3 % |  |  | CD4 % |  |  | CD8 % |  |  |
| --- | --- | --- | --- | --- | --- | --- | --- | --- | --- | --- |
|  |  | M | N | S | M | N | S | M | N | S |
| IL-2/4/7 | IFN-γ median | 4.27 | 4.98 | 10.60 | 4.24 | 5.88 | 10.65 | 2.42 | 4.81 | 0.82 |
| | IFN-γ min | 0.11 | 0.12 | 0.21 | 0.14 | 0.14 | 0.22 | 0.10 | 0.05 | 0.25 |
| | IFN-γ max | 33.60 | 17.10 | 14.80 | 33.40 | 18.90 | 20.10 | 42.00 | 19.40 | 3.99 |
|  | IL-2 median | 0.68 | 1.01 | 0.96 | 0.41 | 0.70 | 0.90 | 0.42 | 0.42 | 0.27 |
|  | IL-2 min | 0.14 | 0.23 | 0.20 | 0.14 | 0.18 | 0.16 | 0.00 | 0.16 | 0.18 |
|  | IL-2 max | 1.41 | 1.85 | 2.31 | 1.44 | 2.09 | 1.82 | 1.20 | 0.71 | 0.79 |
| | TNF-α median | 2.80 | 2.35 | 3.99 | 2.64 | 2.09 | 4.59 | 1.45 | 0.97 | 0.61 |
| | TNF-α min | 0.19 | 0.22 | 0.25 | 0.05 | 0.04 | 0.07 | 0.10 | 0.00 | 0.09 |
| | TNF-α max | 12.90 | 7.50 | 7.86 | 12.90 | 8.41 | 7.90 | 5.65 | 3.70 | 2.42 |
| IL-2/7/15 | IFN-γ median | 6.05 | 3.60 | 4.47 | 6.00 | 3.61 | 4.55 | 0.33 | 3.13 | 1.15 |
| | IFN-γ min | 0.15 | 0.17 | 0.48 | 0.13 | 0.21 | 0.42 | 0.09 | 0.04 | 0.12 |
| | IFN-γ max | 20.80 | 15.50 | 26.00 | 34.20 | 16.10 | 29.40 | 47.30 | 20.50 | 2.65 |
|  | IL-2 median | 1.03 | 0.53 | 0.79 | 0.68 | 0.58 | 0.80 | 0.37 | 0.49 | 0.41 |
|  | IL-2 min | 0.15 | 0.32 | 0.31 | 0.20 | 0.28 | 0.22 | 0.20 | 0.23 | 0.27 |
|  | IL-2 max | 2.55 | 1.56 | 6.59 | 1.64 | 0.91 | 6.52 | 2.24 | 1.16 | 1.75 |
| | TNF-α median | 4.18 | 1.70 | 2.37 | 3.90 | 1.46 | 2.74 | 0.69 | 1.29 | 0.76 |
| | TNF-α min | 0.24 | 0.38 | 0.78 | 0.10 | 0.48 | 0.33 | 0.00 | 0.10 | 0.24 |
| | TNF-α max | 7.06 | 4.20 | 10.10 | 12.60 | 4.50 | 11.20 | 18.50 | 7.27 | 3.61 |
| IL-2/4/21 | IFN-γ median | 0.30 | 0.20 | 0.36 | 0.51 | 0.29 | 0.16 | 0.00 | 0.00 | 0.00 |
| | IFN-γ min | 0.03 | 0.14 | 0.08 | 0.00 | 0.00 | 0.00 | 0.00 | 0.00 | 0.00 |
| | IFN-γ max | 2.46 | 0.64 | 2.53 | 4.99 | 1.08 | 3.63 | 3.37 | 0.22 | 0.97 |
|  | IL-2 median | 0.16 | 0.21 | 0.16 | 0.13 | 0.20 | 0.16 | 0.00 | 0.00 | 0.00 |
|  | IL-2 min | 0.10 | 0.09 | 0.09 | 0.00 | 0.09 | 0.06 | 0.00 | 0.00 | 0.00 |
|  | IL-2 max | 0.40 | 0.84 | 1.02 | 0.52 | 0.33 | 0.81 | 0.24 | 0.48 | 2.17 |
| | TNF-α median | 0.40 | 0.15 | 0.14 | 0.34 | 0.20 | 0.14 | 0.00 | 0.00 | 0.00 |
| | TNF-α min | 0.10 | 0.09 | 0.00 | 0.06 | 0.00 | 0.00 | 0.00 | 0.00 | 0.00 |
| | TNF-α max | 0.93 | 0.47 | 1.66 | 1.95 | 0.50 | 2.36 | 3.70 | 7.69 | 0.00 |
| IL-2/7/21 | IFN-γ median | 0.74 | 0.25 | 0.29 | 1.27 | 0.16 | 0.18 | 0.00 | 0.33 | 0.41 |
| | IFN-γ min | 0.14 | 0.06 | 0.11 | 0.07 | 0.00 | 0.00 | 0.00 | 0.00 | 0.00 |
| | IFN-γ max | 0.97 | 0.38 | 0.50 | 2.93 | 0.81 | 1.36 | 1.52 | 5.82 | 1.11 |
|  | IL-2 median | 0.09 | 0.07 | 0.11 | 0.20 | 0.22 | 0.14 | 0.10 | 0.13 | 0.34 |
|  | IL-2 min | 0.03 | 0.05 | 0.07 | 0.10 | 0.00 | 0.00 | 0.00 | 0.00 | 0.00 |
|  | IL-2 max | 0.73 | 0.66 | 0.31 | 1.27 | 1.10 | 0.57 | 1.75 | 0.60 | 0.83 |
| | TNF-α median | 0.19 | 0.19 | 0.20 | 0.45 | 0.33 | 0.59 | 0.39 | 0.53 | 0.10 |
| | TNF-α min | 0.13 | 0.07 | 0.14 | 0.17 | 0.19 | 0.18 | 0.00 | 0.00 | 0.00 |
| | TNF-α max | 1.70 | 0.31 | 0.26 | 2.55 | 1.01 | 2.58 | 0.94 | 0.72 | 0.93 |

Percent IFN-γ , IL-2 and TNF-α production (median, minimum and maximum values) from the CD3+, CD4+ and CD8+ compartments of SARS-CoV-2 T cells stimulated with the peptide libraries derived from M, N and S structural proteins after expansion under the different culture conditions with the four different cytokine cocktails IL-2/4/7, IL-2/7/15, IL-2/4/21 and IL-2/7/21.

**Figure 1.**
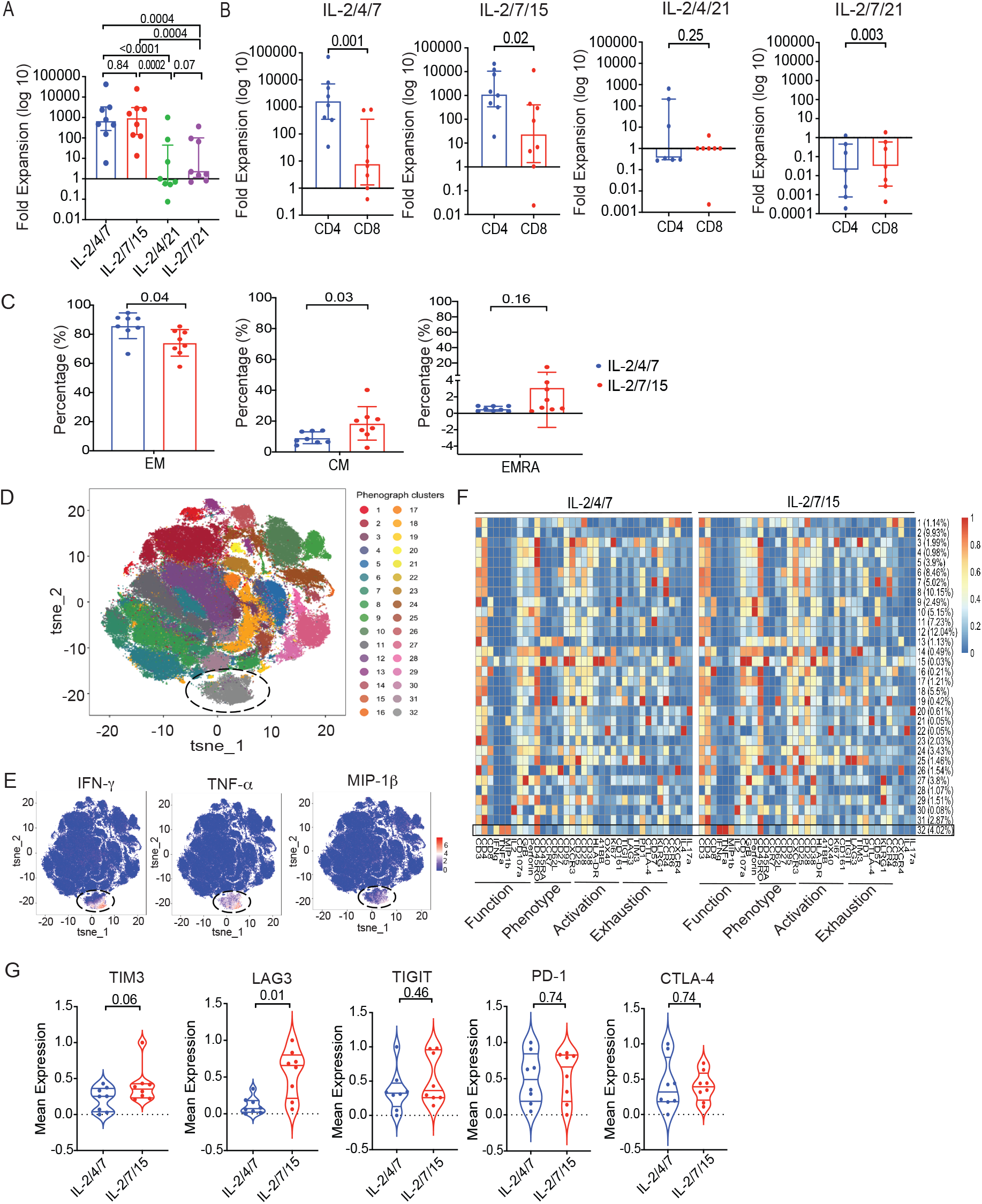
Successful expansion of SARS-CoV-2 T cells from COVID-19 recovered donors. **A**, Bar graph showing the log 10 fold expansion of SARS-CoV-2 T cells cultured with different cytokine cocktails, IL-2/4/7 (blue), IL-2/7/15 (red), IL-2/4/21 (green) and IL-2/7/21 (purple). **B**, Bar graphs showing the log 10 fold expansion of the CD4+ (blue) and CD8+ (red) subsets of SARS-CoV-2 T cells cultured under the different cytokine stimulation conditions. **C**, Quantification of EM (CD45RO+ CD45RA-CD62L-, left panel), CM (CD45RO+ CD45RA-CD62+, middle panel) and EMRA (CD45RO-CD45RA+ CD62L-, right panel) in SARS-CoV-2 T cells expanded with IL-2/4/7 (blue) or IL-2/7/15 (red) (n=8 samples per group); bars represent median values with interquartile range. p-values are indicated at the top of each graph. **D**, Mass cytometry analysis of T cells (gated on CD45+CD3+) expanded from 8 recovered donors using combination of M, N and S peptide libraries and cytokine cocktails (IL-2/4/7 and IL-27/15 conditions are overlapped in this phenograph). tSNE map shows the 32 clusters obtained, each highlighted in corresponding color. Cluster 32 (circled) represents the polyfunctional SARS-CoV-2 T cells. **E**, Individual tSNE maps show the expression of IFN-γ, TNF-α and MIP-1β mostly restricted to cluster 32. Expression levels are indicated by color scale, ranging from blue representing low expression to red representing high expression. **F**, Cluster identity and frequency are summarized in heatmaps showing marker expression levels (X axis) for T cell populations (Y axis) expanded with the two different cytokine cocktails IL-2/4/7 (left heatmap) or IL-2/7/15 (right heatmap). Markers associated with function, phenotype, activation or exhaustion are indicated below each heatmap. Expression level is indicated by color scale, ranging from low (blue) to high (red). Cluster 32 is indicated with a black rectangle. **G**, Violin plots comparing expression of TIM3, LAG3, TIGIT, PD-1, and CTLA-4 and between the two cytokine stimulation conditions IL-2/4/7 (blue) and IL-2/7/15 (red), n = 8. p-values are indicated at the top of each graph.

### SARS-CoV-2 reactive T cells generated from CoV-RD are polyfunctional

We next interrogated the functional phenotype of the *ex vivo* expanded SARS-CoV-2 CTLs. Since IL-2/4/7 and IL-2/7/15 resulted in the best cell expansion, we focused our analysis on SARS-CoV-2 CTLs generated using these two conditions. Both culture conditions supported expansion of effector memory (EM) and central memory (CM) T cells although the use of IL-2/7/15 resulted in expansion of CM T cells (**Figure 1C**).

Previous studies in patients with severe COVID-19 have reported the presence of T cells with an exhausted phenotype and reduced polyfunctionality (Chen et al., 2020; Zheng et al., 2020a, 2020b). Thus, we performed a comprehensive single cell analysis of expanded SARS-CoV-2 CTLs from 8 recovered donors using mass cytometry. Phenotypic interrogation of SARS-CoV-2 reactive T-cells expanded with IL-2/4/7 or IL-2/7/15 (identified based on their ability to produce IFN-γ in response to *ex vivo* stimulation with a mixture of S, M and N peptide libraries) revealed that SARS-CoV-2 specific CTLs are polyfunctional based on their ability to secrete multiple cytokines and chemokines simultaneously, including IFN-γ, TNF-α and MIP-1β (cluster 32; **Figure 1D and 1E**). Moreover, *ex vivo* expanded SARS-CoV-2 CTLs did not express high levels of inhibitory/checkpoint molecules thus, arguing against an exhausted phenotype (cluster 32; **Figure 1F**). Indeed, analysis of functional markers revealed a cytotoxic Th1 phenotype, characterized by expression of IFN-γ, TNF-α, CD107a and granzyme B (GrB), indicating direct antiviral killing capacity (**Figure 1F**). Interestingly, they did not produce significant amounts of IL-2 in response to antigenic stimulation. Single cell phenotypic comparison of SARS-CoV-2 CTLs expanded using the two different culture conditions did not reveal major differences in the expression patterns of activation and functional markers between these two groups (**Figure 1F**). However, cells expanded in the presence of IL-2/4/7 expressed lower levels of some exhaustion markers such as TIM3 and LAG3 compared to cells expanded with IL-2/7/15 (**Figure 1G**). Taken together, these data support the notion that polyfunctional, non-exhausted T cells capable of reacting against SARS-CoV-2 antigens can be expanded from the PB of CoV-RDs.

We also performed a multiplex analysis to measure cytokines in supernatants collected from cultures of SARS-CoV-2 CTLs with SARS-CoV-2 antigens (n= 4 for each of the culture conditions-IL-2/4/7 and IL-2/7/15). As expected, the expanded SARS-CoV-2 CTLs released effector cytokines such as IFN-γ, TNF-α, MIP-1β in response to antigenic stimulation; of note, they did not produce cytokines such as IL-6, IL-1α, or IL-10 that could contribute to a higher risk of toxicity or cytokine release syndrome (CRS) (**Figure S1**).

### Expanded SARS-CoV-2 CTLs from CoV-RD are directed against structural proteins, including both the C and N terminals of the S protein

In order to identify the dominant antigen(s) driving expansion of SARS-CoV-2 CTLs, the expanded cells were stimulated *ex vivo* with peptide libraries derived from either M, N, S or E (structural proteins) or AP3A, Y14, NS6 NS7a, NS7B, NS8, ORF9B or ORF10 (non-structural proteins). Analysis of IFN-γ production showed that for the lines expanded with IL-2/4/7, the overall T-cell response was mostly directed against S (median 10.60%, range 0.21 –14.8%), with the remaining cells responding to M (median 4.27%, range 0.11– 33.6%) or N (median 4.98%, range 0.12 –17.10%) (**Figure 2A**). For the lines expanded with IL-2/7/15, the CD3+ T-cell response favored M (median 6.05%, range 0.15 – 20.80%) followed by S (median 4.47%, range 0.48 – 26.00%) and N (median 3.60 %, range 0.17-15.50%) (**Figure 2A**). In sum, for both IL-2/4/7 and IL-2/7/15 culture settings, seven lines were directed against M + N + S, one line was directed against S + N, and no line reacted to M and N in the absence of S. There was no significant expansion of CTLs in response to the non-structural proteins or the structural E protein.

**Figure 2.**
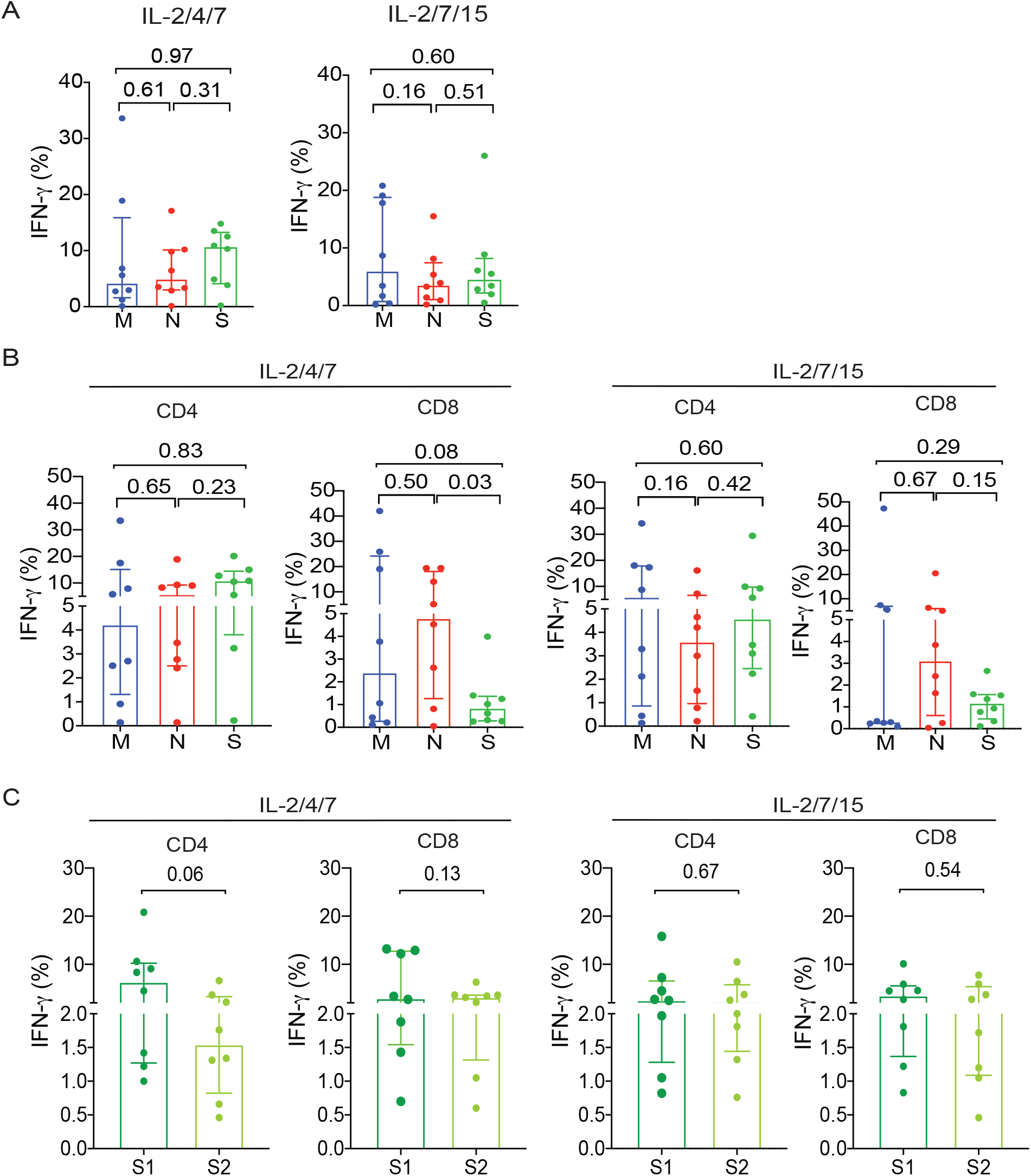
Expanded SARS-CoV-2 CTLs are directed against structural proteins including the C and N terminals of the S protein. **A**, Bar graphs showing the percentage of total IFN-γ (+) SARS-CoV-2 CD3+ T cells stimulated with the peptide libraries derived from the different structural proteins M (blue), N (red), S (green) cultured with different cytokine cocktails IL-2/4/7 (left panel) or IL-2/7/15 (right panel). **B**, Bar graphs showing IFN-γ (+) expression in CD4+ and CD8+ T cell subsets of SARS-CoV-2 T cells stimulated with the peptide libraries derived from the different structural proteins M (blue), N (red), S (green) cultured with IL-2/4/7 (left panels) or IL-2/7/15 (right panels). **C**, Quantification of IFN-γ (+) SARS-CoV-2 T cells subsets (CD4+ or CD8+) directed against N-terminus (S1, dark green) or the C-terminus (S2, light green) of the S protein in both IL-2/4/7 (left panels) and IL-2/7/15 (right panels) stimulation conditions (n=8 samples per group). Bars represent median values with interquartile range. p-values are indicated at the top of each graph.

When we considered the CD4+ and CD8+ T cell responses separately, we found that the response of CD4+ T cells to individual SARS-CoV-2 antigens followed a pattern similar to that observed in the overall CD3+ T cell population. Interestingly, however, CD8+ T cell responses were mostly directed against the N protein, irrespective of the cytokine cocktail used for SARS-CoV-2 CTL expansion (**Figure 2B, Table 1**).

Peptides derived from the C-terminus of the S protein have higher homology with the S glycoprotein of human endemic “common cold” coronaviruses; in contrast, the N-terminus of the S protein includes peptides from the receptor-binding domain (the target of neutralizing antibodies) that are more specific to SARS-CoV-2 (Braun et al., 2020; Walls et al., 2020). Our expanded CD4+ and CD8+ SARS-CoV-2 CTLs were capable of reacting to both N- and C-terminal epitopes (Pools 1 and 2 of the S protein respectively), indicating their specificity for the receptor binding domain (RBD) of SARS-CoV-2 (**Figure 2C, Tables 2, S4 and S5**).

**Table 2.** Cytokine production of SARS-CoV-2 T cells from recovered donors expanded with different cytokine cocktails against S1 and S2 (N and C terminals of the S protein).

|  |  | CD3 % |  | CD4 % |  | CD8 % |  |
| --- | --- | --- | --- | --- | --- | --- | --- |
|  |  | S1 | S2 | S1 | S2 | S1 | S2 |
| **IL-2/4/7** | IFN-γ median | 5.81 | 3.88 | 6.40 | 1.55 | 3.02 | 3.26 |
| | IFN-γ min | 1.33 | 0.71 | 1.00 | 0.46 | 0.70 | 0.60 |
| | IFN-γ max | 26.50 | 8.54 | 20.80 | 6.60 | 13.20 | 6.29 |
|  | IL-2 median | 0.53 | 0.51 | 0.41 | 0.38 | 1.78 | 2.87 |
|  | IL-2 min | 0.14 | 0.24 | 0.06 | 0.07 | 0.00 | 0.15 |
|  | IL-2 max | 5.03 | 4.48 | 2.55 | 2.72 | 6.15 | 6.47 |
| | TNF-α median | 3.24 | 1.41 | 2.56 | 0.87 | 5.51 | 1.80 |
| | TNF-α min | 0.45 | 0.72 | 0.33 | 0.51 | 0.00 | 0.00 |
| | TNF-α max | 9.47 | 2.63 | 12.80 | 1.60 | 27.30 | 14.70 |
| **IL-2/7/15** | IFN-γ median | 5.94 | 2.99 | 2.61 | 2.27 | 3.59 | 2.22 |
| | IFN-γ min | 1.73 | 1.61 | 0.82 | 0.76 | 0.83 | 0.46 |
| | IFN-γ max | 18.00 | 14.60 | 15.80 | 10.50 | 10.10 | 7.77 |
|  | IL-2 median | 0.80 | 0.90 | 0.22 | 0.27 | 3.03 | 1.87 |
|  | IL-2 min | 0.12 | 0.05 | 0.05 | 0.03 | 0.14 | 0.40 |
|  | IL-2 max | 4.37 | 4.97 | 3.02 | 2.34 | 4.13 | 4.48 |
| | TNF-α median | 2.20 | 2.02 | 2.62 | 1.36 | 10.27 | 11.38 |
| | TNF-α min | 1.57 | 0.95 | 0.84 | 0.38 | 0.36 | 0.00 |
| | TNF-α max | 10.50 | 4.14 | 13.90 | 3.05 | 26.20 | 22.60 |
| **IL-2/4/21** | IFN-γ median | 0.17 | 0.14 | 0.14 | 0.22 | 0.00 | 0.00 |
| | IFN-γ min | 0.03 | 0.08 | 0.00 | 0.05 | 0.00 | 0.00 |
| | IFN-γ max | 4.15 | 0.26 | 6.21 | 0.60 | 0.67 | 0.00 |
|  | IL-2 median | 0.17 | 0.16 | 0.14 | 0.16 | 0.00 | 0.00 |
|  | IL-2 min | 0.00 | 0.08 | 0.00 | 0.00 | 0.00 | 0.00 |
|  | IL-2 max | 0.40 | 0.57 | 0.68 | 0.44 | 0.66 | 0.61 |
| | TNF-α median | 0.23 | 0.14 | 0.29 | 0.18 | 0.00 | 0.00 |
| | TNF-α min | 0.07 | 0.05 | 0.08 | 0.00 | 0.00 | 0.00 |
| | TNF-α max | 1.78 | 0.72 | 2.69 | 0.30 | 0.00 | 1.59 |
| **IL-2/7/21** | IFN-γ median | 0.20 | 0.12 | 0.23 | 0.10 | 0.43 | 0.00 |
| | IFN-γ min | 0.05 | 0.05 | 0.00 | 0.00 | 0.00 | 0.00 |
| | IFN-γ max | 0.89 | 0.63 | 2.22 | 1.24 | 0.63 | 0.47 |
|  | IL-2 median | 0.08 | 0.08 | 0.17 | 0.39 | 0.00 | 0.35 |
|  | IL-2 min | 0.06 | 0.04 | 0.09 | 0.11 | 0.00 | 0.00 |
|  | IL-2 max | 0.11 | 0.21 | 0.37 | 1.50 | 0.54 | 0.87 |
| | TNF-α median | 0.20 | 0.09 | 0.35 | 0.21 | 0.00 | 0.00 |
| | TNF-α min | 0.08 | 0.06 | 0.20 | 0.07 | 0.00 | 0.00 |
| | TNF-α max | 0.37 | 0.31 | 1.18 | 1.50 | 0.63 | 0.83 |

Percent IFN-γ , IL-2 and TNF-α production (median, minimum and maximum values) from the CD3+, CD4+ and CD8+ compartments of SARS-CoV-2 T cells stimulated with the peptide libraries derived from S1 and S2 (N and C terminals of the S protein) after expansion under different cytokine stimulation conditions (IL-2/4/7, IL-2/7/15, IL-2/4/21 and IL-2/7/21).

### Culture with IL-2/4/7 results in preferential expansion of T cells against the N-terminus of the S protein

Since the choice of cytokines used to expand SARS-CoV-2 T-cells modulates the hierarchy of antigenic response, we compared the T cell response against S (S1 and S2), M and N proteins of SARS-CoV-2 in PB samples at baseline (prior to expansion) with that observed in paired expanded CTLs from Cov-RDs.

At baseline (pre-expansion), the responses were mostly CD4 dominant (**Figure 3A**), and directed against the S protein (median 0.21%, range 0.02%-0.56%), followed by N (median 0.15%, range 0.01-0.33%) and M (median 0.11%, range 0.01-0.24%) **(Figure 3B**), indicating an immunogenic dominance for S protein HLA class II epitopes. We did not detect measurable responses against non-structural proteins (data not shown). Following expansion, culture with IL-2/4/7 maintained the hierarchy of CD4+ T cell response toward S protein, (**Figures 2B, 3C, and 3D**), with a greater proportion of T cells directed against S1, while culture with IL-2/7/15 favored a response toward M protein (**Figures 2B, 3C, and 3D**). Expansion with IL-2/4/21 and IL-2/7/21 yielded very low numbers of CTLs, most of which were CD4+. Interestingly, the pattern of antigenic response for cells cultured using these two conditions differed from that observed for IL-2/4/7 and was similar to IL-2/7/15, in that the majority of CD4+ CTLs reacted instead to M and N proteins (**Figures S2A**). For CTLs that did show a response to S protein, further analysis detected no particular pattern of reactivity to either terminus of the protein (**Figure S2B**).

**Figure 3.**
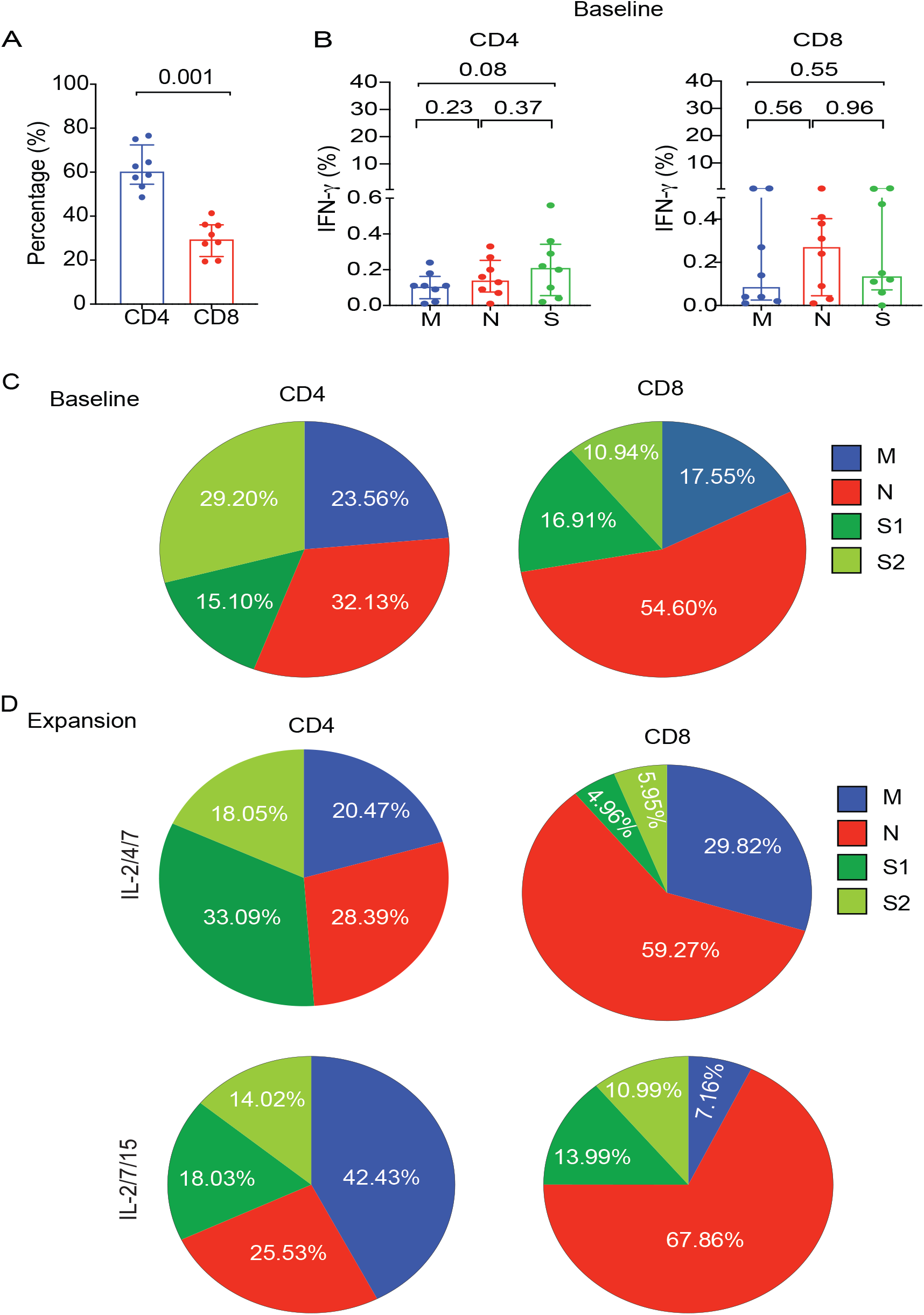
Pattern of antigenic responses after expansion of SARS-CoV-2 T cells from COVID-19 recovered donors compared to baseline. **A**, Bar graph showing the percentage of CD4+ and CD8+ subsets of SARS-CoV-2 T cells at baseline. **B**, Graphical analysis showing the percentage of IFN-γ (+) SARS-CoV-2 T cells from recovered donors at baseline in the CD4+ compartment (left panel) or CD8+ compartment (right panel) when stimulated with the peptide libraries derived from the different structural proteins M (blue), N (red), S (green). **C**,**D** Pie charts illustrating the percent distribution of IFN-γ (+) CD4 and CD8 T cells reactive to M (blue), N (red) or S1 (dark green), S2 (light green) peptide libraries at baseline **(C)** or following expansion with IL-2/4/7 (upper panel) or IL-2/7/15 (lower panel) cytokine cocktails **(D)**.

Furthermore, we found a significant correlation (p=0.002, R^2^=0.82) between the spike protein IgG antibody titer measured in plasma from the recovered donors and the absolute number of SARS-CoV-2 specific T cells following expansion with IL-2/4/7, but not with the IL-2/7/15 cytokine cocktail (**Figure S3**).

These data indicate that the antigenic skewing can be driven both by the immunodominance of the protein and by the culture conditions and support the use of IL-2/4/7 for the expansion of SARS-CoV-2 T-cells for clinical use.

### SARS-CoV-2 T-cells can be expanded from the PB of healthy donors but at lower frequencies than for Cov-RD donors

Recent reports indicate the presence of SARS-CoV-2 T cells in the PB of healthy donors (HD) not exposed to COVID-19 (Braun et al., 2020; Pia, 2020). Thus, we asked if SARS-CoV-2 T-cells can be expanded from the PB of HD and whether they have a similar pattern of SARS-CoV-2 recognition as those generated from Cov-RDs. PBMC from 5 HDs were expanded using the same protocol as for Cov-RDs. We achieved only a modest expansion of CTLs recognizing SARS-CoV-2 antigens over a 14-day culture period, with a median 20.37-fold increase (range, 2.85 – 41.84) for IL-2/4/7 culture condition and 21.49-fold increase (range, 4.00 – 53.95) for IL-2/7/15 culture condition (**Figure 4A)**. The frequencies of SARS-CoV-2 CTLs from the PB of healthy donors after 14 days of culture were significantly lower than those achieved with PB from Cov-RDs (**Figure 4B**). At baseline (pre-expansion), assessment of IFN-γ production and CTL frequencies suggested that in healthy donors responses were directed mostly against the N protein (median 0.13%, range 0.03%-0.37%), followed by S (median 0.10%, range 0.04-0.12%) and M (median 0.09%, range 0.01-0.75%) (**Figure S4A**). Culture in the presence of either cytokine cocktail could skew the response towards S, albeit at much lower frequencies than that observed with Cov-RDs (**Figure S4B, Tables S6 and S7**). We did not find any particular pattern of antigenic response in the CD4 and CD8 compartments (**Figure S4B, Tables S6 and S7**). Expansion was not successful in IL-2/4/21 or IL-2/7/21 stimulation conditions.

**Figure 4.**
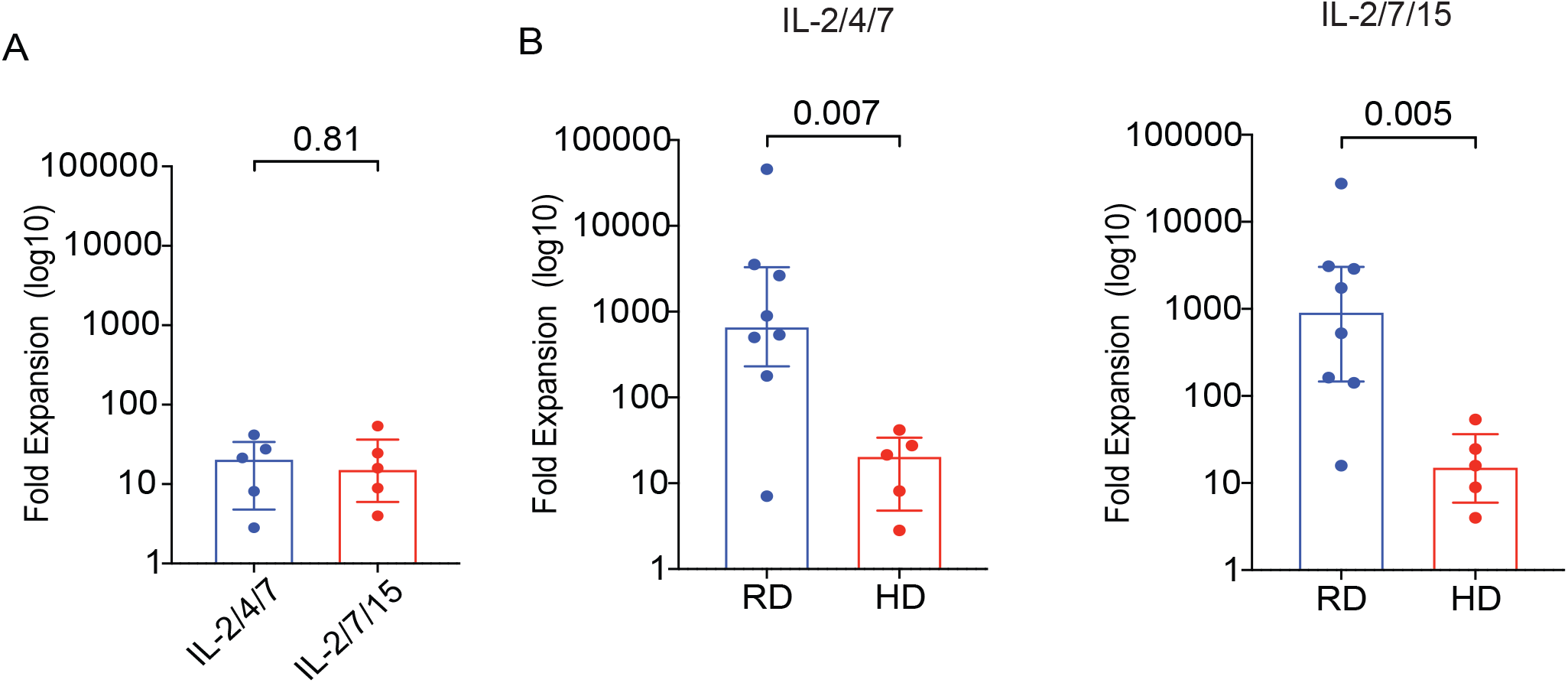
SARS-CoV-2 CTLs can be expanded from the PB of healthy donors but at lower frequencies compared to recovered donors. **A**, Graphical representation of the log 10 fold expansion of SARS-CoV-2 T cells derived from healthy donors cultured with different cytokine cocktails, IL-2/4/7 (blue), IL-2/7/15 (red). **B**, Comparison of SARS-CoV-2 T cell expansion between recovered donors (RD, blue) and healthy donors (HD, red) cultured under the different cytokine stimulation conditions IL-2/4/7 (left panel) and IL-2/7/15 (right panel).

### Expanded SARS-CoV-2 T cells can be genetically modified to render them steroid-resistant

Corticosteroids are used in the treatment of patients with COVID-19-related ARDS to reduce mortality associated with this condition. SARS-CoV-2 specific T-cell therapy is not an option in such patients as corticosteroids induce apoptosis of adoptively transferred T cells, thus, significantly limiting the efficacy of this approach. To address this challenge, we used CRISPR/Cas9 gene editing to knockout the glucocorticoid receptor gene *(Nuclear Receptor Subfamily 3 Group C Member 1 [NR3C1]*) in SARS-CoV-2 CTLs and confirmed high efficiency of deletion (>90%) as determined by PCR and western blot analysis (**Figures 5A and 5B**). Annexin V apoptosis assay confirmed that the viability of *NR3C1* KO CTLs treated with dexamethasone was similar to that of control CTLs (defined as CTLs electroporated with Cas9 alone) (**Figures 5C and 5D**). Moreover, *NR3C1* KO SARS-CoV-2 CTLs maintained similar phenotype and distribution of CD4+ and CD8+ T cell subsets when compared to control SARS-CoV-2 CTLs and retained their effector functions (**Figures 5E-5G**).

**Figure 5.**
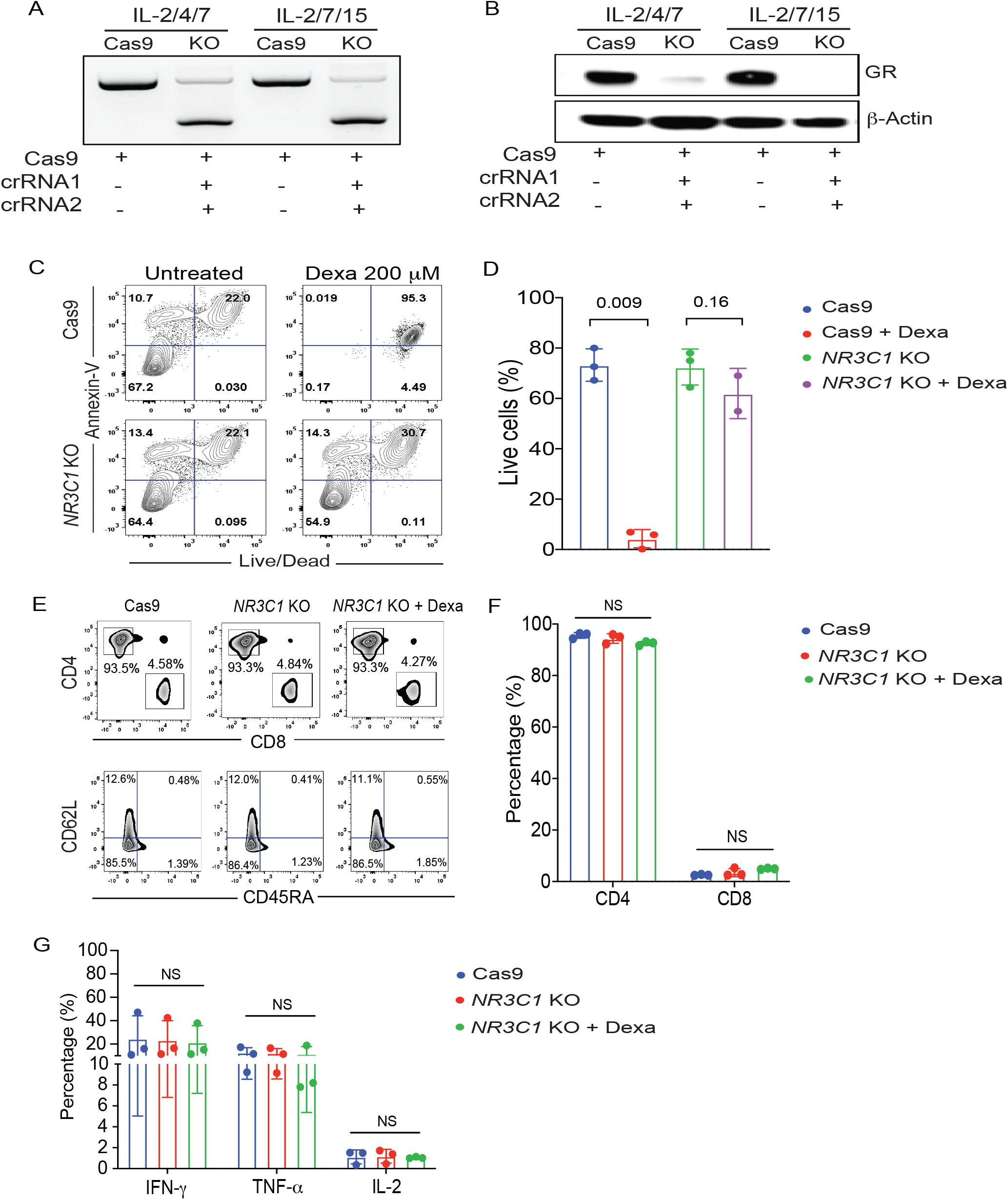
Expanded SARS-CoV-2 CTLs can be genetically modified to become steroid resistant. **A**,**B** *NR3C1* KO efficiency shown by PCR gel electrophoresis **(A)** and by western blot **(B)** in SARS-CoV-2 CTLs expanded with IL-2/4/7 or IL-2/7/15, after electroporation with Cas9 alone (Cas9) or Cas9 complexed with crRNA 1 and crRNA 2 targeting exon 2 of the *NR3C1* gene. SARS-CoV-2 CTLs electroporated with Cas9 alone were used as controls. β-actin was used as loading control for western blot. **C**, Representative FACS plots showing the percentage of apoptotic cells (Annexin V+) and live or dead cells (live/dead stain) in control Cas9 vs *NR3C1* KO SARS-CoV-2 CTLs after culture with or without dexamethasone (Dexa; 200 μM) for 72 hours. Inset values indicate the percentage of annexin V and alive/dead cells from each group. **D**, Bar graph summarizing the percentage of live cells between control Cas9 and *NR3C1* KO SARS-CoV-2 CTLs treated with or without 200 μM dexamethasone for 72 hours (n = 3). Bars represent median values with interquartile range. p-values are indicated above the graphs. **E**, Representative FACs plots showing the distribution of CD4+ and CD8+ T cells (upper panel) and phenotype based on CD62L and CD45RA expression (lower panel) in Cas9 alone or NR3C1 KO SARS-CoV-2 CTLs with or without 200 μM dexamethasone. **F**, Percentage of CD4+ and CD8+ T cells within SARS-CoV-2 CTLs treated with control Cas9 (blue), *NR3C1* KO (red), or *NR3C1* KO plus dexamethasone (Dexa; 200μM; green). **G**, Frequency of SARS-CoV-2 CTLs producing IFN-γ, TNF-α, or IL-2 control Cas9 (blue), *NR3C1* KO (red), or *NR3C1* KO plus dexamethasone (Dexa; 200μM; green) in response to 6 hours of stimulation with viral PepMix (n = 3). The functional analysis of the Cas9 + Dexa group was not performed due to the absence of viable cells resulting from the lymphocytotoxic effect of steroids. The bars represent mean values with SD. NS, not significant.

## DISCUSSION

Here we show that large numbers of SARS-CoV-2 T-cells can be generated from buffy coats of convalescent patients with specificity directed against multiple structural proteins of this virus, including the RBD of the S protein. These cells can be genetically modified to render them resistant to the lymphocytotoxic effect of corticosteroids, thus, making their application clinically feasible.

We performed single cell analysis of the *ex vivo* expanded SARS-CoV-2 CTLs to better understand their phenotypic and functional properties. SARS-CoV-2 T-cells were classified based on their state of differentiation into naïve, central memory, effector memory or terminally differentiated effector memory (TEMRA). SARS-CoV-2 T-cells comprised mostly of effector and central memory T cells. This phenotype predicts for the capacity to persist and provide long-term immunity after adoptive transfer (Powell et al., 2005). We also investigated their functional state based on their ability to produce one or more cytokines in response to *ex vivo* stimulation with SARS-CoV-2 antigens. Polyfunctionality is defined as the production of multiple cytokines by T cells and is associated with protective immune responses to viruses and vaccines (Minton, 2014). We confirmed that SARS-CoV-2 T-cells were polyfunctional and predominantly of Th1 phenotype.

T cells derived from patients with severe COVID-19 have been reported to express multiple inhibitory molecules (Song et al., 2020), raising concerns that following *ex vivo* expansion, they may have an exhausted phenotype with poor effector function and replicative senescence. In our study, T cells from COVID-19-recovered individuals did not have an exhaustion phenotypic signature following *in vitro* expansion, and retained their functional phenotype. This observation is consistent with reversal of exhausted phenotype upon antigen clearance, a phenomenon also reported in other virus infection settings (Wieland et al., 2017).

SARS-CoV-2 neutralizing antibodies are directed against the RBD within the N-terminal of the S glycoprotein. Peptides derived from this region of the protein are believed to be specific to SARS-CoV-2, while peptides from the C terminus are shared with other beta-coronaviruses (Braun et al., 2020). We showed that expanded SARS-CoV-2 T-cells from CoV-RD were capable of recognizing epitopes from both the N- and C-terminus of the S glycoprotein, indicating specificity for the SARS-CoV-2 virus. In addition, expanded CTLs reacted against N and M proteins, which are reportedly also shared among different beta-coronaviruses (Patrick et al., 2006). The specific antigens that drive an effective and protective T-cell response against SARS-CoV-2 are not yet known. They may be proteins that are shared with other beta-coronaviruses, or they may be unique to SARS-CoV-2 (e.g. RBD) or may likely be a combination of both.

Cytokines can modulate the phenotype of T cells by activating different signaling pathways. We tested different cytokine cocktails to identify the optimal conditions for promoting the expansion of SARS-CoV-2 T-cells with a memory phenotype and without evidence of exhaustion. Since both CD4 and CD8 T cells are involved in successful antiviral response, we also investigated whether cytokines would support expansion of both subsets. We observed that cocktails including IL-2/4/7 or IL-2/7/15 resulted in expansion of clinically relevant doses of polyfunctional SARS-CoV-2 T-cells with a central and effector memory phenotype. Interestingly, the combination of IL-2/4/7 preferentially supported expansion of T cells against S and in particular the RBD-containing S1 region of the protein, making this the cytokine cocktail of choice for the production of SARS-CoV-2 T-cells for clinical use. Of note, the inclusion of IL-21 in the cytokine cocktail resulted in poor expansion of SARS-CoV-2 T-cells, as previously observed in other memory T cell expansion studies (Li et al., 2005).

CRS is a major complication of COVID-19 (Moore and June, 2020) and is caused by the production of inflammatory cytokines such as IL-6 by virus-infected myeloid cells. The inflammatory milieu overstimulates cells of the innate and adaptive immune system that in turn contribute to the observed cytokine storm (Kang et al., 2019). Therefore, a legitimate concern with our approach is that the adoptively infused SARS-CoV-2 T-cells could amplify CRS and worsen the patient’s condition. However, we believe that ACT for COVID-19 is unlikely to worsen CRS as the adoptively infused CTLs will target and kill the SARS-CoV-2 infected myeloid cells, thus breaking the vicious cycle driving the cytokine storm. Furthermore, CRS is not unique to beta-coronavirus infections and has been reported with other viral infections such as CMV, EBV and adenovirus (Humar et al., 1999; McLaughlin et al., 2018; Ramos-Casals et al., 2014) where adoptive cell therapy with virus-specific CTLs has been used to treat hundreds of patients with severe infections effectively and with minimal complications (Bollard and Heslop, 2016; McLaughlin et al., 2018; Muftuoglu et al., 2018; Tzannou et al., 2017).

Our approach allows for cryopreservation and banking of SARS-CoV-2 T-cells, facilitating the rapid identification and selection of viral-specific T-cells for ACT based on the most closely HLA-matched third-party donor as published by our group and others for other severe viral infections (Eiz-Vesper et al., 2012; Haque et al., 2007; Leen et al., 2011; Muftuoglu et al., 2018; O’Reilly et al., 2016). An additional advantage to our approach is that the genetic modification of SARS-CoV-2 T-cells to delete the glucocorticoid receptor will allow treatment of patients with ARDS on high doses corticosteroids.

In summary, adoptive transfer of SARS-CoV-2 T-cells may be a suitable therapeutic strategy for treatment of patients with severe COVID-19. We intend to initiate a clinical study at MD Anderson Cancer Center to test this approach in patients in the near future.

## ACKNOWLEDGEMENTS

Supported in part by the generous philanthropic contributions to The University of Texas MD Anderson Cancer Center AML Moonshot Program, by the grants from National Institute of Health, National Cancer Institute 5R01CA211044-04 and Cancer Center Support (CORE) Grant (CA016672) that support the Flow Cytometry and Cellular Imaging Facility and the RNA sequencing core facility at MD Anderson Cancer Center.

## AUTHOR CONTRIBUTIONS

Conceptualization, R.B., K.R.; Methodology, R.B., N.U, E.E., M.D.; Investigation, N.U, E.E., M.S., B.H., E.G., M.M., F.R.S., S.A., S.L., L.M.G., P.L.; Writing, K.R., R.B., N.U., E.E., M.D.,T.L., L.F.-M., D.M.; Funding Acquisition, K.R.; Resources K.K., F.M., F.A., H.S., S.G.Y., Q.M., J.D., K.C., D.M., A.L.; Supervision K.R., P.P.B., R.C., E.J.S.

## STAR METHODS

### Key Resources Tables

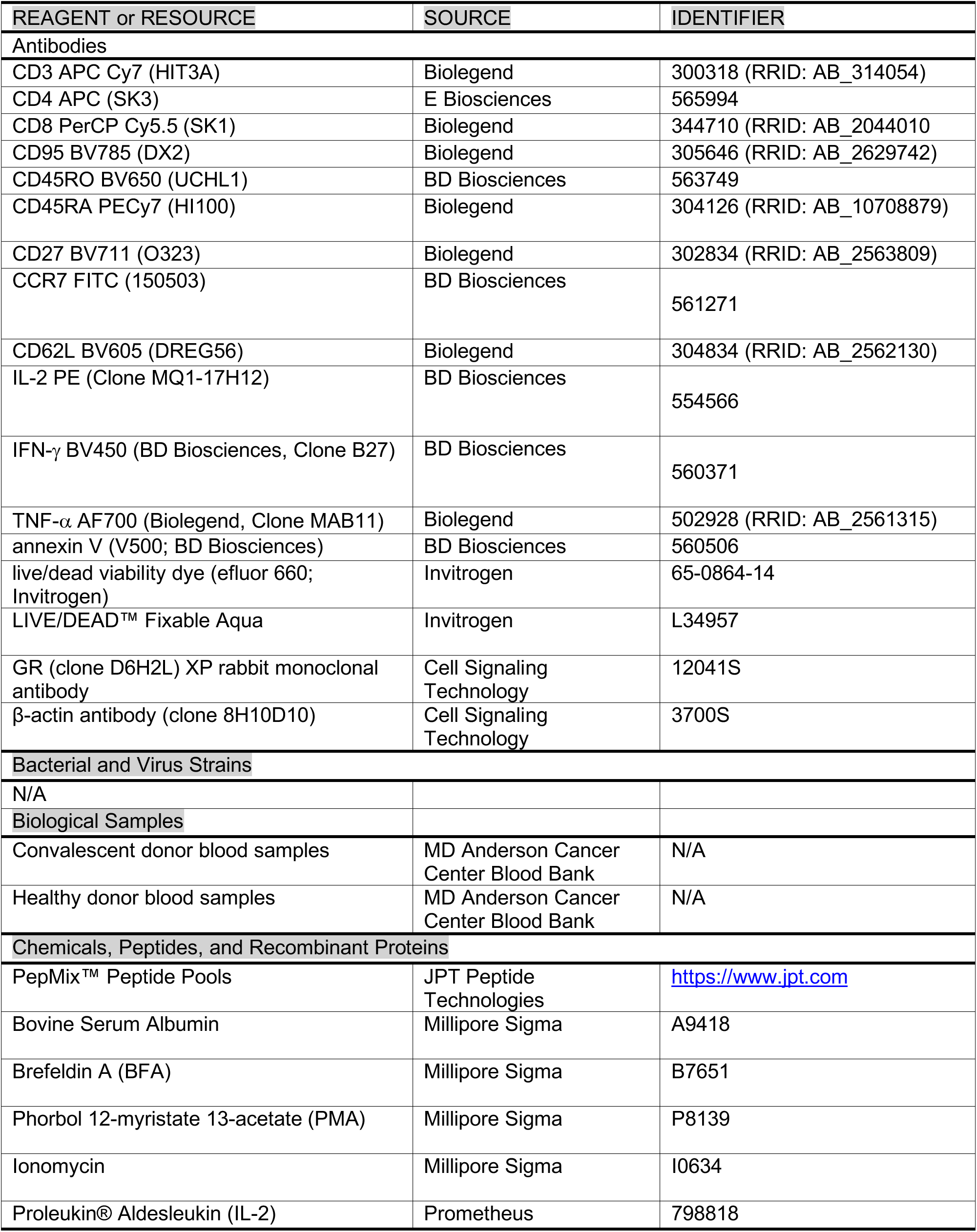
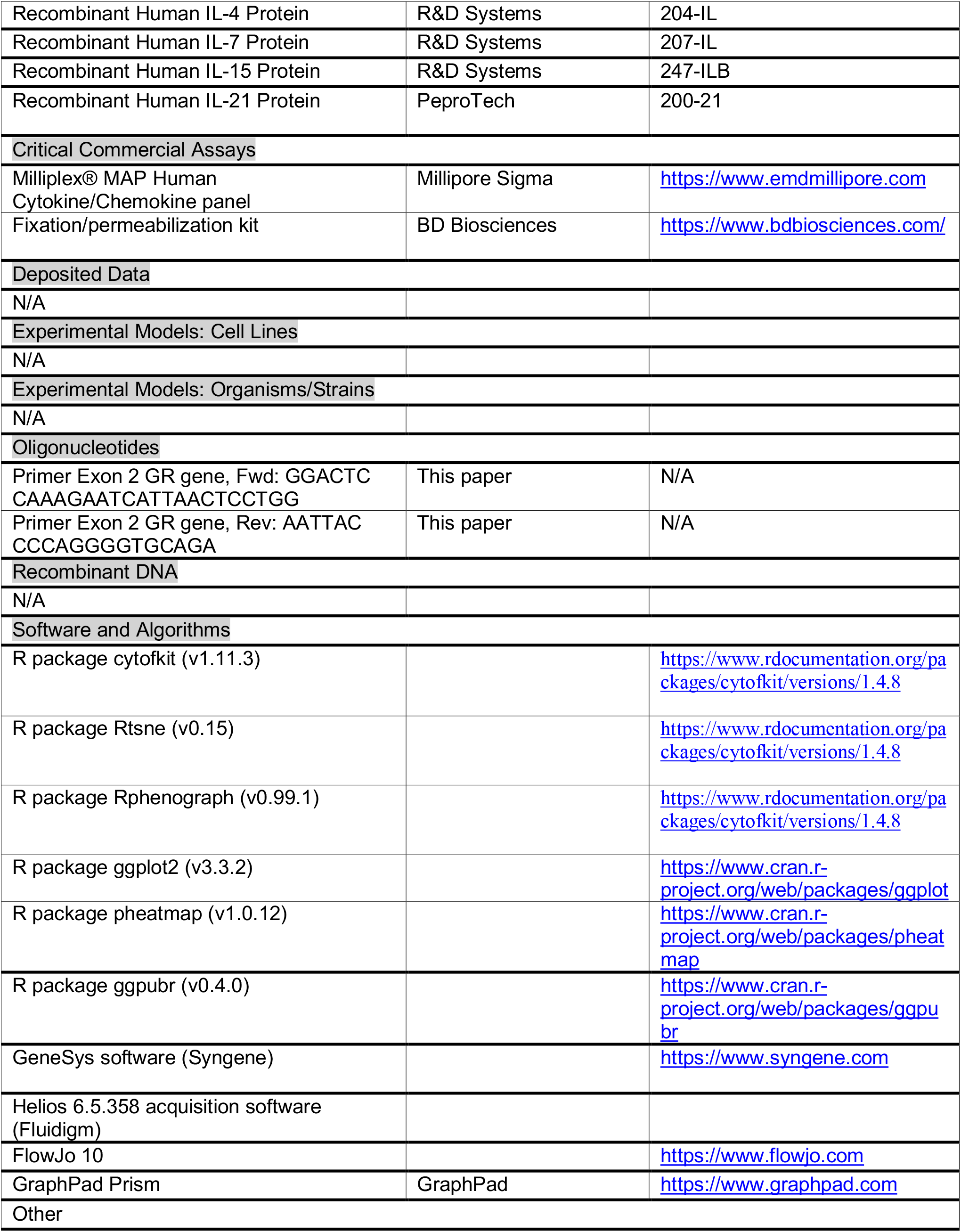

### Resource Availability

#### Lead Contact

Further information and requests may be directed to and will be fulfilled by the Lead Contact Katayoun Rezvani

### Materials Availability

All requests for data and materials will be reviewed by MD Anderson Cancer Center to verify if the request is subject to any intellectual property or confidentiality obligations. Any data and materials that can be shared by the corresponding author will be released freely or via a Material Transfer Agreement if deemed necessary.

### Data and Code Availability

No unique code was generated.

### Experimental Model and Subject Details

#### COVID-19 recovered donors and healthy donors

Buffy coat units were processed from 500mL of whole blood collected from each of the 10 COVID-19 recovered donors (CoV-RD) and 20 mL of peripheral blood from 5 healthy donors were collected under local Institutional Review Board approved protocols (Lab02-0630 and PA13-0647) and following informed consent. All donors were 18 years or older and were recruited without consideration of disease severity, race, ethnicity or gender. All CoV-RD had recovered from proven symptomatic COVID-19 confirmed by a positive test for SARS-CoV-2. At the time of blood collection, all were asymptomatic for at least 14 days and had a negative PCR test, confirming full recovery.

Blood from Cov-RD was collected in heparin-coated blood bags and stored at room temperature prior to processing for peripheral blood mononuclear cell (PBMC) isolation. PBMCs were isolated by density-gradient sedimentation using Ficoll-Paque (Lymphoprep, Oslo, Norway). Isolated PBMCs were either used fresh for *ex vivo* expansion of SARS-CoV-2 specific T cells (SARS-CoV-2 CTLs) or cryopreserved in freezing media containing 10% DMSO (GIBCO), supplemented with 10% heat inactivated Human Serum AB (Gemini Bio) and stored in liquid nitrogen until used for phenotypic and functional assays.

### Functional assessment of SARS-CoV-2 reactive T cells

For intracellular assessment of cytokine production, cells were stimulated *ex vivo* with 15mer pepMixes overlapping by 11 amino acids derived from SARS-CoV-2 spike (S) (peptide pool 1 or 2), membrane (M), nucleocapsid (N), envelope (E), or the non-structural proteins (AP3A, Y14, NS6, NS7a, NS7B, NS8,ORF9B and ORF10) (JPT, Germany) [1µg /ml per peptide] for 4 hours. Stimulation with an equimolar amount of DMSO was performed as negative control and with Phorbol 12-myristate 13-acetate (PMA)-Ionomycin (1.25ng/ul and 0.05ng/μl, respectively) as positive control. Brefeldin A (Millipore Sigma, St. Louis, MO) was added into the culture for 4 hours. Cells were stained with an antibody cocktail containing live/dead viability dye Aqua (Invitrogen), CD3 APC Cy7 (Biolegend, Clone HIT3A), CD4 APC (E Biosciences, Clone SK3), CD8 PerCP Cy5.5 (Biolegend, Clone SK1), CD95 BV785 (Biolegend, Clone DX2), CD45RO BV650 (BD Biosciences, Clone UCHL1), CD45RA PECy7 (Biolegend, Clone HI100), CD27 BV711 (Biolegend, Clone O323), CCR7 FITC (BD Biosciences, Clone 150503) and CD62L BV605 (Biolegend, Clone DREG56) for 30 minutes on ice, then fixed and permeabilized using the BD fixation/permeabilization kit (BD Biosciences, San Diego, CA) according to manufacturer’s protocol. Cells were subsequently stained with antibodies against IL-2 PE (BD Biosciences, Clone MQ1-17H12), IFN-γ BV450 (BD Biosciences, Clone B27), and TNF-α AF700 (Biolegend, Clone MAB11) for 30 mins. Following a final wash, cells were re-suspended in FACS buffer and data were acquired on a BD LSRFortessa (BD Biosciences). Data analysis was performed using Flowjo (Tree Star, Ashland, OR). The gates applied for the identification of IFN-γ, IL-2, and TNF-α on the total population of CD4+ and CD8+ T-cells were defined according to the negative control for each individual. Similar functional assays were performed for *NR3C1* knockout (KO) CTLs.

### SARS-CoV-2 antibody assay

IgM and IgG responses against nucleocapsid, S1 receptor-binding domain (RBD), S1S2, S2, S1, OC43, HKU1, NL63 Nucleoprotein, and 229E Spike derived from SARS-CoV-2 and other human coronaviruses were performed at Genalyte (Austin, TX) CLIA-certified laboratory using plasma from convalescent patients.

### Cytokine and chemokine measurement

Cells were stimulated *ex vivo* with 15mer pepMixes from S, M and N for 24 hours at 37°C and 5% CO2. Supernatants were collected and assayed with the Milliplex® MAP Human Cytokine/Chemokine panel (EMD Millipore Corporation, Burlington, MA) following the manufacturer’s instructions.

### Generation of SARS-CoV-2 specific T-cells

Isolated PBMC from CoV-RD and HD were pulsed with a SARS-CoV-2 pepMix (JPT, Germany) comprising the entire length of the structural (S, M, N, E) and non-structural (AP3A, Y14, NS6, NS7a, NS7B, NS8, ORF9B and ORF10) proteins at a concentration of 1 µg/ml per peptide. Cells were cultured in complete media with 5% human AB serum and supplemented with four different cytokine cocktails: IL-2 (50 IU/ml), IL-4 (60 ng/ml) and IL-7 (10 ng/ml) vs. IL-2 (50 IU/ml), IL-7 (10 ng/ml) and IL-15 (10 ng/ml) vs. IL-2 (50 IU/ml), IL-4 (60 ng/ml) and IL-21 (30 ng/ml) vs. IL-2 (50 IU/ml), IL-7 (10 ng/ml) and IL-21 (30 ng/ml) every 3 days. After 14 days of expansion, the frequencies of SARS-CoV-2 specific T-cells were determined by intracellular cytokine staining.

### Mass Cytometry

A panel of 40 metal-tagged antibodies was used for the in-depth characterization of SARS-CoV-2 reactive T-cells (**Table S1)**. All unlabeled antibodies were purchased in carrier-free form (Fluidigm) and conjugated in-house with the corresponding metal tag using Maxpar X8 polymer per the manufacturer’s instructions (Fluidigm) and as previously described (Muftuoglu et al., 2018). Briefly, thawed PBMCs were rested overnight at 37°C / 5% CO2 and stained with a freshly prepared antibody mix against cell surface markers for 30 minutes at room temperature on a shaker (100 rpm). For the last 3 minutes of incubation, cells were incubated with 2.5 µM cisplatin (Pt198, Fluidigm) for viability assessment, washed twice with cell staining buffer and fixed/permeabilized using BD fixation/permeabilization solution for 30 minutes in dark at 4°C. Cells were washed twice with perm/wash buffer, stained with antibodies directed against intracellular markers and after an additional wash step, stored overnight in 500 µl of 1.6% paraformaldehyde (EMD Biosciences)/PBS with 125 nM iridium nucleic acid intercalator (Fluidigm). Samples were supplemented with EQ calibration beads (Fluidigm) and acquired at 300 events/second on a Helios instrument (Fluidigm) using the Helios 6.5.358 acquisition software (Fluidigm). Mass cytometry data were normalized based on EQ™ four element signal shift over time using Fluidigm normalization software 2. Initial data processing was performed using Flowjo version 10.2. Calibration beads were gated out and singlets were chosen based on iridium 193 staining and event length. Dead cells were excluded by the Pt198 channel and manual gating was performed to select the CD45+CD3+ population which was subsequently exported for downstream analyses. A total of 156,384 cells were evenly sampled from 16 samples derived from 8 patients to perform automated clustering analysis. The data were processed using the R package cytofkit (v1.11.3). Expression values for each marker were arcsine transformed with a cofactor of 5. Data dimensionality reduction was performed using the R package Rtsne (v0.15) for t-Distributed Neighbor Embedding (tSNE) analysis. The R package Rphenograph (v0.99.1) was used to cluster all cells into 32 clusters. Both the R package Rstne (v0.15) and the R package Rphenograph (v0.99.1) were implemented in the R package cytofkit (v1.11.3). The t-SNE plots were generated using the R package ggplot2 (v3.3.2). Normalized mean values of marker expressions in each cluster were plotted as heatmap using the function “pheatmap” from R package pheatmap (v1.0.12). Min-max normalization was used to scale each marker’s mean expressions range to [0,1]. The normalized mean values of marker expressions were plotted as box plots using the function “ggpaired” from R package ggpubr (v0.4.0). The mean comparison p-values of Wilcoxon signed-rank test were added to the plots using the function “stat_compare_means” from R package ggpubr (v0.4.0).

### CRISPR-Cas9 gene editing of the glucocorticoid receptor

Knockout (KO) of *NR3C1* (the glucocorticoid receptor gene) was performed on day 7 of T cell expansion using ribonucleoprotein (RNP) complex. We used two crRNAs targeting exon 2 of the human *NR3C1* gene: crRNA #1 TGAGAAGCGACAGCCAGTGA, crRNA#2 GGCCAGACTGGCACCAACGG as previously described.(Basar et al., 2020) Briefly, Cas9 protein (IDT) and gRNA (crRNA + tracrRNA combination) were complexed and electroporated into 1 million SARS-CoV-2 specific T cells using the Neon transfection system (Thermo Fisher Scientific).

### Annexin V apoptosis assay

Annexin V apoptosis assay was performed to evaluate the effect of dexamethasone on the viability of CTLs from Cas9 control and *NR3C1* KO groups. CTLs from both groups were treated with 200 µM dexamethasone (Sigma) for 72 hours. Cells were then collected, washed with annexin V buffer, and stained with annexin V (V500; BD Biosciences) and live/dead viability dye (efluor 660; Invitrogen) in addition to CD3 APC Cy7 (Biolegend, Clone HIT3A), CD4 APC (E Biosciences, Clone SK3), and CD8 PerCP Cy5.5 (Biolegend, Clone SK1). The proportion of apoptotic (positive for annexin V) and dead CTLs (positive for live/dead stain) was determined by flow cytometry.

### PCR gel electrophoresis

DNA was extracted and purified (QIAamp DNA Blood Mini Kit; Qiagen Inc) from SARS-CoV-2 specific T cells (control and *NR3C1* KO conditions). We used the Platinum SuperFi Green PCR Master Mix from Invitrogen for polymerase chain reaction (PCR) amplification using the following PCR primers spanning the Cas9–single-guide RNA cleavage site of exon 2 of the GR gene: exon 2 forward primer, GGACTCCAAAGAATCATTAACTCC TGG; exon 2 reverse primer, AATTACCCCAGGGGTGCAGA. DNA bands were separated by agarose gel electrophoresis prepared with SYBR-safe DNA gel stain in 0.5× Tris/Borate/EDTA. Gel images were obtained using GeneSys software in a G:BOX gel documentation system (Syngene).

### Western blot

To detect GR protein expression, CTLs were lysed in lysis buffer (IP Lysis Buffer; Pierce Biotechnology Inc) supplemented with protease inhibitors (Complete Mini, EDTA-free Cocktail tablets; Roche Holding) and incubated for 30 minutes on ice. Protein concentration was determined by the bicinchoninic acid (BCA) assay (Pierce Biotechnology Inc). The following primary antibodies were used: GR (clone D6H2L) XP rabbit monoclonal antibody and β-actin antibody (clone 8H10D10); both antibodies were obtained from Cell Signaling Technology. Blots were imaged using a G:BOX gel documentation system and GeneSys software (Syngene).

