## Supplementary Materials for "Generation of glucocorticoid resistant SARS-CoV-2 T-cells for adoptive cell therapy"

### Supplemental Information

### Supplemental Figures

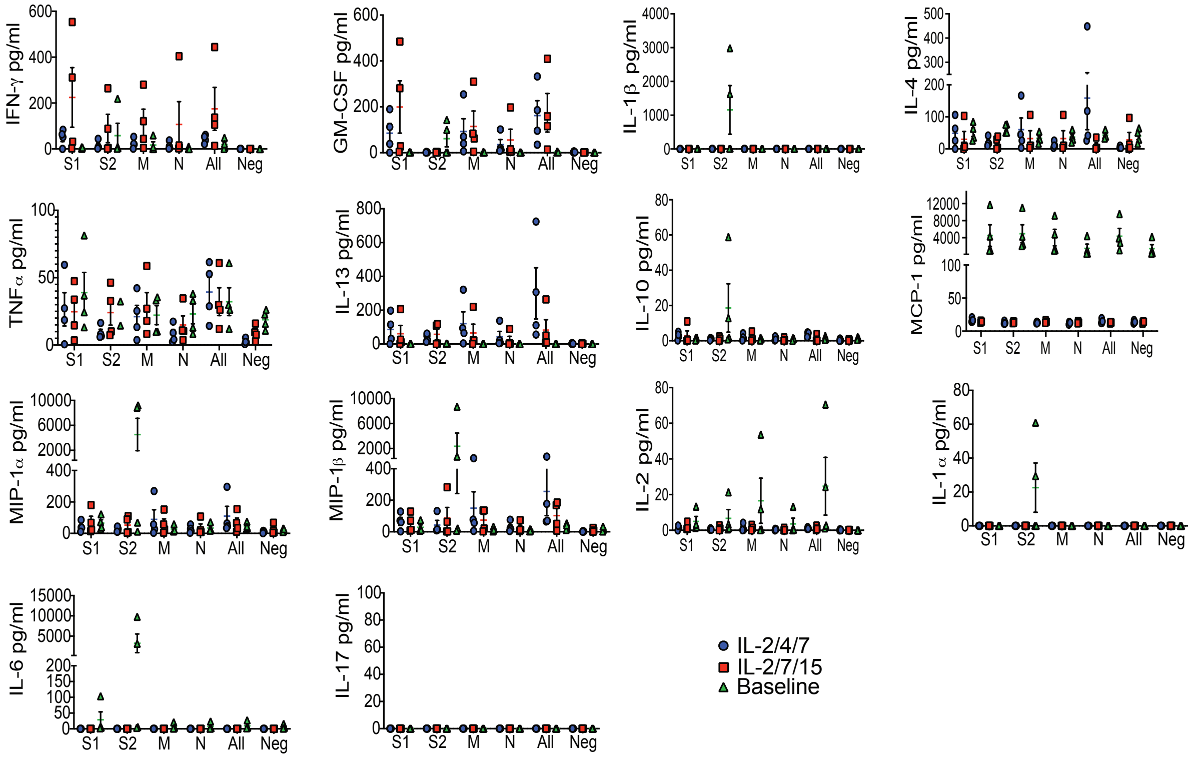


**Figure S1. Cytokine profile from COVID-19 reactive T cells supports a functional profile without CRS, related to Figure 1.** Multiplex cytokine analysis showing the concentration of different cytokines in pg/ml detected in supernatants from COVID-19 reactive T cells stimulated with the different peptide libraries (S1, S2, M and N, either separately or in combination) at baseline (green triangles) or after expansion with IL-2/4/7 (blue circles) or IL-2/7/15 (red squares). n=4 samples per group. Neg refers to negative control without peptide stimulation.

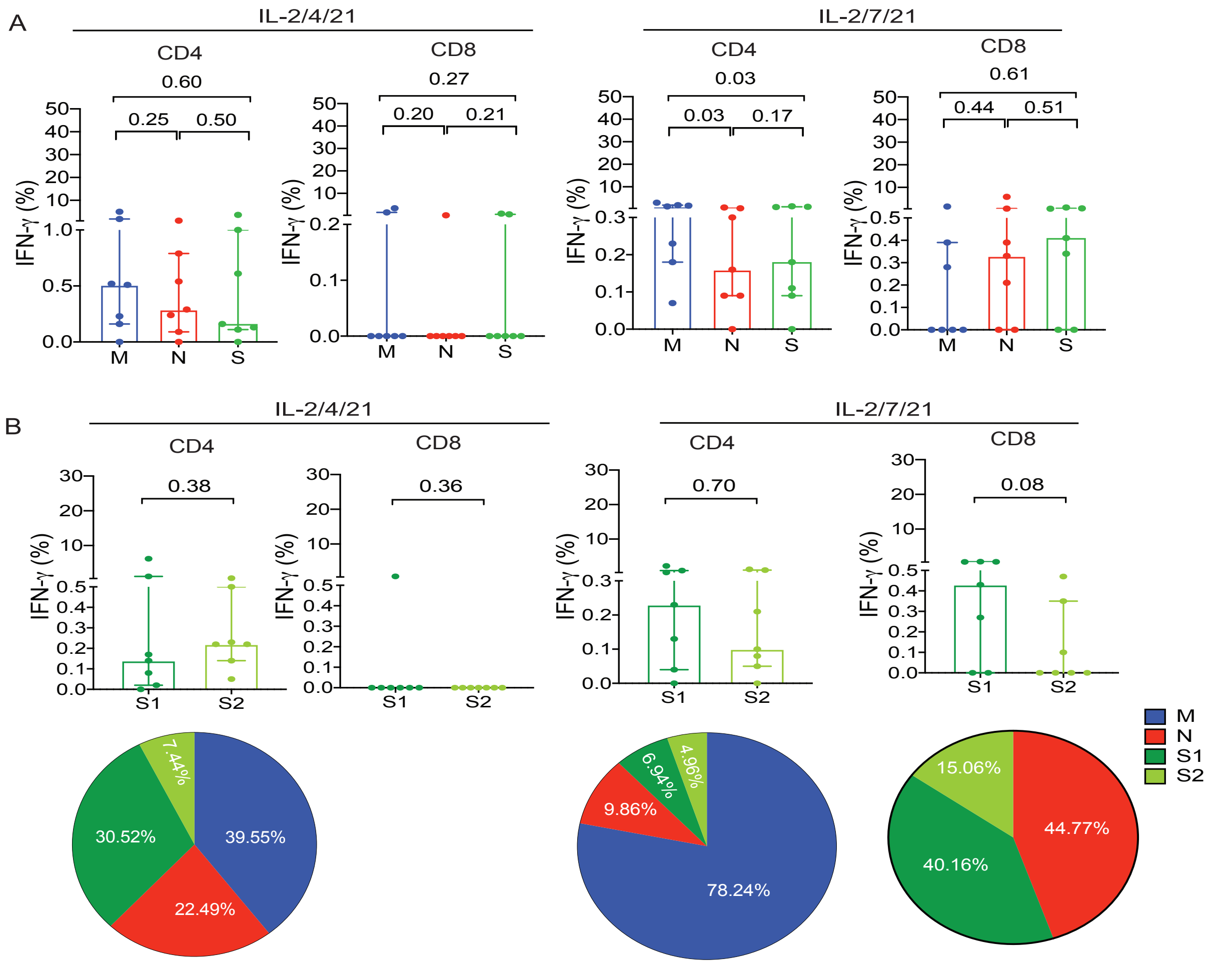


**Figure S2. Expanded COVID-19 CTLs are directed against structural proteins, including both the C and N terminals of the S protein, related to Figure 3. A**, Percentage of IFN-γ (+) COVID-19 reactive CD3+ T cells stimulated with the peptide libraries derived from the different structural proteins M (blue), N (red), S (green) cultured with different cytokine cocktails IL2/4/21 (left panel) or IL2/7/21 (right panel). **B,** Quantification of IFN-γ (+) COVID-19 reactive T cells subsets (CD4+ or CD8+) directed against N-terminus (S1, dark green) or the C-terminus (S2, light green) of the S protein in both IL-2/4/21 (left panels) and IL-2/7/21 (right panels) stimulation conditions (n=7 samples per group). Bars represent median values with interquartile range. p-values are indicated at the top of each graph. Corresponding pie charts showing the percent distribution of M (blue), N (red), S1 (dark green) and S2 (light green) reactive IFN-γ (+) T cells are depicted under each bar graph. Please note that a pie chart was not generated for the CD8+ T cells cultured with IL-2/4/21 due to the very low number of cells. Inset percentages (%) within each pie chart represent the fraction of IFN-γ (+) T cells that are reactive to specific peptides. Sum of all fractions is 100% representing the total IFN-γ (+) T cells.

A

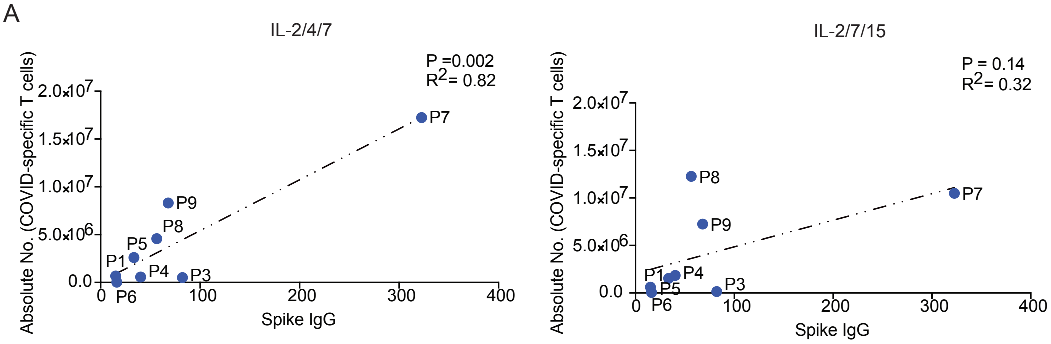


**Figure S3. Absolute number of COVID-19 specific T cells cultured with IL-2/4/7 correlates with antibody titer of Spike protein IgG, related to Figure 3.** Scatter plots showing correlation between absolute number of COVID-19 specific T cells (on Y axis) and antibody titers on the X axis (Spike IgG) of COVID-19 specific T cells expanded with IL-2/4/7 (left panel) or IL-2/7/15 (right panel).

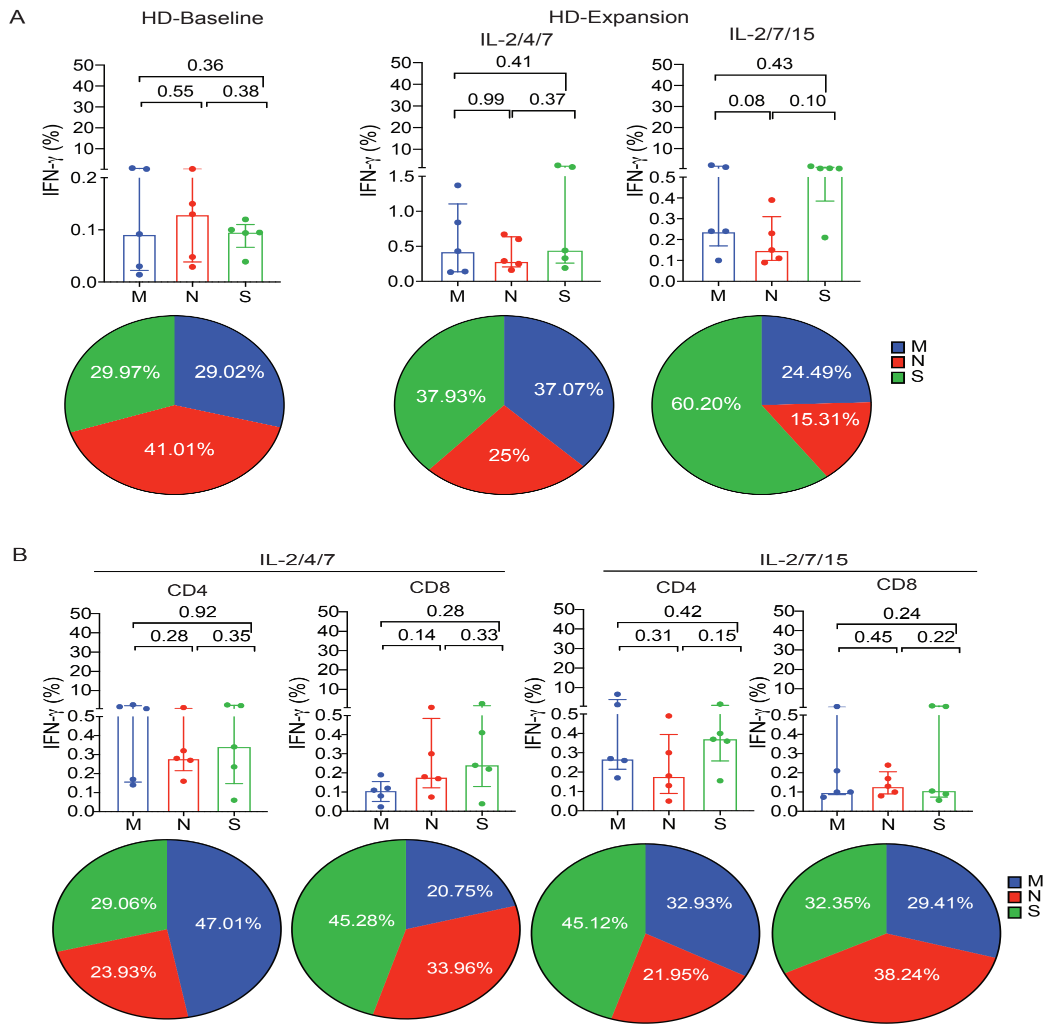


**Figure S4. COVID-19 CTLs can be expanded from the PB of healthy donors but at lower frequencies compared to COVID-19-recovered donors, related to Figure 4. A, B** Graphical analysis of IFN-γ (+) COVID-19 reactive CD3+ T cells from healthy donors stimulated with the peptide libraries derived from the different structural proteins M (blue), N (red), S (green) at **(A)** baseline or **(B)** cultured with IL-2/4/7 (left panel) or IL-2/7/15 (right panel). Corresponding pie charts show the distribution of M (blue), N (red) or S (green) reactive IFN-γ (+) T cells for each group (n=5 samples per group). Bars represent median values with interquartile range. p-values are indicated at the top of each graph. **C,** Percentage of IFN-γ (+) COVID-19 reactive CD4 and CD8 T cells generated from PB of healthy donors stimulated with the peptide libraries derived from the different structural proteins M (blue), N (red), S (green) after expansion with IL-2/4/7 (left panels) or IL-2/7/15 (right panels). Distribution of M (blue), N (red) or S (green) reactive IFN-γ (+) T cells for each group is shown in pie charts below corresponding graphs (n=5 samples per group). Bars represent median values with interquartile range. p-values are indicated at the top of each graph. Inset percentages (%) within each pie chart represent the fraction of IFN-γ (+) T cells that are reactive to specific peptides. Sum of all fractions is 100% representing the total IFN-γ (+) T cells.

### Supplemental Tables

**Table S1. Mass cytometry panel used for phenotyping of COVID-19 reactive T cells, related to Figure 1.**

| Tag | Antibody | Clone | Source |
|---|---|---|---|
| 89Y | CD45 | HI30 | Fluidigm |
| 141Pr | CCR6 | G034E3 | Fluidigm |
| 142Nd | IL4 | MP425D2 | Fluidigm |
| 143Nd | CD4 | OKT4 | Biolegend |
| 144Nd | IL2 | MQ1-17H12 | BD Biosciences |
| 145Nd | CD62L | DREG-56 | Biolegend |
| 146Nd | CD8 | RPA-T8 | Biolegend |
| 147Sm | CD127 | MB15-18C9 | Miltenyi Biotec |
| 148Nd | IL17a | BL168 | Biolegend |
| 149Sm | CD25 | 2A3 | BD Biosciences |
| 150Nd | CD28 | CD28.2 | BD Biosciences |
| 151Eu | CD107a | H4A3 | BD Biosciences |
| 152Sm | CD95 | DX2 | BD Biosciences |
| 153Eu | CCR4 | 205410 | R&D Systems |
| 154Sm | TIGIT | MSA43 | Thermo Fisher |
| 155Gd | CD27 | M-T271 | BD Biosciences |
| 156Gd | CXCR4 | 12G5 | Biolegend |
| 158Gd | OX40 | ACT35 | Biolegend |
| 159Tb | Perforin | delta G9 | Miltenyi Biotec |
| 160Gd | CD45RA | HI100 | Biolegend |
| 161Dy | TIM3 | REA635 | Miltenyi Biotec |
| 162Dy | Ki-67 | Ki-67 | Biolegend |
| 163Dy | CXCR3 | G025H7 | Biolegend |
| 164Dy | CD45RO | UCHL1 | Biolegend |
| 165Ho | CTLA-4 | L3D10 | Biolegend |
| 166Er | MIP1b | W15138A | Biolegend |
| 167Er | CCR7 | G043H7 | Biolegend |
| 168Er | IFNg | B27 | Biolegend |
| 169Tm | TNFa | MAb11 | Biolegend |
| 170Er | HLA-DR | L243 | Biolegend |
| 171Yb | CD161 | 191B8 | Miltenyi |
| 172Yb | KLRG1 | 13F12F2 | Thermo Fisher |
| 173Yb | Granzyme B | GB11 | Fluidigm |
| 174Yb | PD1 | PD1.3.1.3 | Miltenyi Biotec |
| 175Lu | LAG3 | 11C3C65 | Biolegend |
| 176Yb | CD38 | REA572 | Miltenyi Biotec |
| 209Bi | 41BB | 4B4-1 | Biolegend |
| 115In | CD57 | HCD57 | Biolegend |
| Pt194 | CD3 | UCTH1 | Biolegend |
| Pt198 | LD | | |

**Table S2. Cytokine production and fold expansion of COVID-19 reactive T cells derived from the different recovered donors and expanded with IL-2/4/7 or IL-2/7/15 cytokine cocktails, related to Figure 1.**

| | | | IL-2/4/7 | | | | | | | | |
|---|---|---|---|---|---|---|---|---|---|---|---|
| | | | CD3 % | | | CD4 % | | | CD8 % | | |
| | Fold Expansion | | M | N | S | M | N | S | M | N | S |
| Patient 1 | 177.37 | IFN-γ | 5.59 | 6.44 | 14.80 | 7.90 | 8.29 | 20.10 | 0.43 | 2.62 | 1.40 |
| | | IL-2 | 0.92 | 0.91 | 2.31 | 0.45 | 0.47 | 1.82 | 0.81 | 0.71 | 0.79 |
| | | TNF-α | 3.18 | 2.67 | 6.52 | 3.14 | 3.10 | 7.12 | 1.45 | 1.13 | 2.03 |
| Patient 3 | 506.42 | IFN-γ | 1.22 | 17.10 | 13.50 | 0.91 | 18.90 | 15.00 | 3.77 | 0.81 | 1.25 |
| | | IL-2 | 0.14 | 1.85 | 1.07 | 0.14 | 2.09 | 1.21 | 0.09 | 0.34 | 0.34 |
| | | TNF-α | 0.67 | 7.50 | 3.99 | 0.57 | 8.41 | 4.59 | 0.66 | 0.00 | 0.11 |
| Patient 4 | 540.69 | IFN-γ | 2.74 | 10.20 | 3.84 | 2.51 | 9.01 | 3.24 | 0.21 | 5.08 | 0.32 |
| | | IL-2 | 0.43 | 1.40 | 0.94 | 0.36 | 1.09 | 0.80 | 0.17 | 0.41 | 0.32 |
| | | TNF-α | 1.66 | 5.35 | 3.23 | 1.63 | 5.06 | 3.26 | 1.05 | 3.01 | 2.42 |
| Patient 5 | 897.59 | IFN-γ | 2.94 | 2.87 | 10.90 | 2.70 | 2.41 | 10.50 | 1.06 | 19.40 | 0.61 |
| | | IL-2 | 0.21 | 0.32 | 0.54 | 0.23 | 0.31 | 0.53 | 0.00 | 0.24 | 0.20 |
| | | TNF-α | 1.23 | 1.18 | 4.42 | 1.30 | 1.26 | 4.64 | 0.53 | 3.07 | 0.61 |
| Patient 6 | 7.16 | IFN-γ | 0.11 | 0.12 | 0.21 | 0.14 | 0.14 | 0.22 | 0.10 | 0.05 | 0.25 |
| | | IL-2 | 0.21 | 0.23 | 0.20 | 0.16 | 0.18 | 0.16 | 0.23 | 0.16 | 0.22 |
| | | TNF-α | 0.19 | 0.22 | 0.25 | 0.05 | 0.04 | 0.07 | 0.10 | 0.12 | 0.32 |
| Patient 7 | 45572.50 | IFN-γ | 33.60 | 9.81 | 10.30 | 33.40 | 9.31 | 10.80 | 42.00 | 19.40 | 3.99 |
| | | IL-2 | 1.41 | 1.10 | 0.97 | 1.44 | 1.11 | 0.99 | 0.61 | 0.43 | 0.22 |
| | | TNF-α | 12.90 | 5.36 | 5.43 | 12.90 | 5.20 | 5.53 | 4.36 | 3.70 | 0.45 |
| Patient 8 | 2645.58 | IFN-γ | 6.81 | 3.33 | 4.87 | 5.77 | 3.46 | 5.48 | 25.90 | 4.53 | 0.27 |
| | | IL-2 | 0.92 | 0.52 | 0.67 | 0.58 | 0.35 | 0.40 | 1.20 | 0.57 | 0.68 |
| | | TNF-α | 2.80 | 0.87 | 1.71 | 2.64 | 0.88 | 1.81 | 5.65 | 0.97 | 0.68 |
| Patient 9 | 3537.23 | IFN-γ | 18.90 | 3.51 | 12.50 | 17.50 | 2.77 | 12.70 | 19.00 | 14.00 | 1.02 |
| | | IL-2 | 3.80 | 1.16 | 1.34 | 3.51 | 0.93 | 1.32 | 1.13 | 1.12 | 0.18 |
| | | TNF-α | 12.50 | 2.35 | 7.86 | 12.20 | 2.09 | 7.90 | 1.83 | 0.56 | 0.09 |

| | | | IL-2/7/15 | | | | | | | | |
|---|---|---|---|---|---|---|---|---|---|---|---|
| | | | CD3 % | | | CD4 % | | | CD8 % | | |
| | Fold Expansion | | M | N | S | M | N | S | M | N | S |
| Patient 1 | 163.62 | IFN-γ | 1.65 | 3.92 | 1.96 | 2.12 | 4.65 | 2.24 | 0.24 | 1.63 | 2.65 |
| | | IL-2 | 1.08 | 1.56 | 1.64 | 0.49 | 0.81 | 0.92 | 0.38 | 0.23 | 1.75 |
| | | TNF-α | 4.18 | 4.20 | 3.98 | 4.22 | 4.50 | 3.67 | 0.38 | 1.29 | 3.61 |
| Patient 3 | 141.91 | IFN-γ | 0.37 | 1.40 | 2.83 | 0.44 | 1.51 | 3.10 | 0.35 | 0.26 | 0.12 |
| | | IL-2 | 0.15 | 0.47 | 0.45 | 0.20 | 0.53 | 0.51 | 1.27 | 0.90 | 1.19 |
| | | TNF-α | 0.39 | 1.97 | 2.37 | 0.52 | 2.22 | 2.74 | 0.00 | 0.52 | 0.24 |
| Patient 4 | 1746.55 | IFN-γ | 8.67 | 8.12 | 26.00 | 8.70 | 4.21 | 29.40 | 0.27 | 20.50 | 0.77 |
| | | IL-2 | 1.24 | 1.31 | 6.59 | 0.87 | 0.91 | 6.52 | 0.37 | 0.76 | 0.29 |
| | | TNF-α | 3.93 | 3.60 | 10.10 | 3.99 | 2.41 | 11.20 | 0.69 | 7.27 | 0.76 |
| Patient 5 | 530.27 | IFN-γ | 3.43 | 3.28 | 6.08 | 3.29 | 3.00 | 5.64 | 0.30 | 2.41 | 1.53 |
| | | IL-2 | 0.35 | 0.33 | 0.45 | 0.34 | 0.30 | 0.43 | 0.20 | 0.38 | 0.41 |
| | | TNF-α | 1.40 | 1.40 | 2.31 | 1.52 | 1.46 | 2.47 | 0.91 | 1.25 | 0.82 |
| Patient 6 | 15.97 | IFN-γ | 0.15 | 0.17 | 0.48 | 0.13 | 0.21 | 0.42 | 0.09 | 0.04 | 0.34 |
| | | IL-2 | 0.26 | 0.32 | 0.31 | 0.22 | 0.28 | 0.22 | 0.52 | 0.74 | 0.58 |
| | | TNF-α | 0.24 | 0.48 | 0.78 | 0.10 | 0.60 | 0.33 | 0.09 | 0.10 | 0.28 |
| Patient 7 | 27716.61 | IFN-γ | 20.80 | 5.36 | 5.51 | 34.20 | 7.19 | 9.23 | 7.35 | 3.84 | 1.58 |
| | | IL-2 | 0.98 | 0.59 | 0.65 | 1.64 | 0.85 | 0.95 | 0.26 | 0.26 | 0.27 |
| | | TNF-α | 7.06 | 2.83 | 3.33 | 12.60 | 4.19 | 5.89 | 1.27 | 1.47 | 0.33 |
| Patient 8 | 2912.07 | IFN-γ | 19.10 | 15.50 | 3.43 | 17.30 | 16.10 | 3.46 | 47.30 | 5.02 | 0.93 |
| | | IL-2 | 2.55 | 1.04 | 0.92 | 0.94 | 0.62 | 0.69 | 2.24 | 1.16 | 0.36 |
| | | TNF-α | 4.90 | 1.70 | 1.91 | 3.90 | 1.43 | 1.82 | 18.50 | 1.45 | 1.14 |
| Patient 9 | 3088.38 | IFN-γ | 17.80 | 0.90 | 8.86 | 18.00 | 0.78 | 9.89 | 5.45 | 6.26 | 1.36 |
| | | IL-2 | 2.12 | 0.40 | 0.93 | 2.08 | 0.36 | 0.91 | 0.33 | 0.49 | 0.45 |
| | | TNF-α | 5.14 | 0.38 | 3.53 | 5.41 | 0.48 | 3.84 | 0.33 | 0.38 | 0.65 |

Percent IFN-γ, IL-2 and TNF-α production from total CD3+, and CD4+ and CD8+ subsets of COVID-19 reactive T cells derived from each recovered donor and stimulated with the peptide libraries derived from M, N and S structural proteins and fold expansion after culture with IL-2/4/7 or IL-2/7/15 cytokine cocktails.

**Table S3. Cytokine production and fold expansion of COVID-19 reactive T cells derived from the different recovered donors and expanded with IL-2/4/21 or IL-2/7/21 cytokine cocktails, related to Figure 1.**

| | | | IL-2/4/21 | | | | | | | | |
|---|---|---|---|---|---|---|---|---|---|---|---|
| | | | CD3 % | | | CD4 % | | | CD8 % | | |
| | Fold Expansion | | M | N | S | M | N | S | M | N | S |
| Patient 3 | 0.57 | IFN-γ | 0.03 | 0.14 | 0.09 | 0.00 | 0.09 | 0.16 | 0.00 | 0.00 | 0.00 |
| | | IL-2 | 0.12 | 0.09 | 0.09 | 0.13 | 0.09 | 0.16 | 0.00 | 0.00 | 0.00 |
| | | TNF-α | 0.10 | 0.14 | 0.14 | 0.06 | 0.00 | 0.12 | 0.00 | 0.00 | 0.00 |
| Patient 4 | 0.53 | IFN-γ | 0.30 | 0.52 | 0.36 | 0.52 | 0.79 | 0.13 | 0.00 | 0.00 | 0.00 |
| | | IL-2 | 0.30 | 0.26 | 0.82 | 0.52 | 0.20 | 0.63 | 0.00 | 0.00 | 0.00 |
| | | TNF-α | 0.40 | 0.17 | 0.21 | 0.26 | 0.00 | 0.13 | 0.00 | 7.69 | 0.00 |
| Patient 5 | 82.96 | IFN-γ | 1.11 | 0.20 | 2.53 | 1.81 | 0.29 | 3.63 | 0.00 | 0.00 | 0.00 |
| | | IL-2 | 0.12 | 0.13 | 0.13 | 0.13 | 0.09 | 0.15 | 0.00 | 0.00 | 2.17 |
| | | TNF-α | 0.66 | 0.09 | 1.66 | 0.87 | 0.13 | 2.36 | 0.00 | 0.00 | 0.00 |
| Patient 6 | 0.08 | IFN-γ | 0.30 | 0.19 | 1.15 | 0.16 | 0.00 | 0.61 | 0.00 | 0.00 | 0.00 |
| | | IL-2 | 0.40 | 0.84 | 1.02 | 0.31 | 0.30 | 0.81 | 0.00 | 0.00 | 0.00 |
| | | TNF-α | 0.90 | 0.47 | 0.00 | 0.47 | 0.45 | 0.00 | 3.70 | 0.00 | 0.00 |
| Patient 7 | 996.18 | IFN-γ | 2.46 | 0.64 | 0.55 | 4.99 | 1.08 | 1.00 | 3.37 | 0.22 | 0.63 |
| | | IL-2 | 0.10 | 0.10 | 0.15 | 0.11 | 0.10 | 0.06 | 0.24 | 0.00 | 0.00 |
| | | TNF-α | 0.93 | 0.34 | 0.28 | 1.95 | 0.50 | 0.42 | 0.72 | 0.22 | 0.00 |
| Patient 8 | 0.71 | IFN-γ | 0.07 | 0.27 | 0.08 | 0.23 | 0.54 | 0.00 | 0.00 | 0.00 | 0.00 |
| | | IL-2 | 0.26 | 0.25 | 0.29 | 0.00 | 0.33 | 0.14 | 0.00 | 0.48 | 0.56 |
| | | TNF-α | 0.19 | 0.13 | 0.12 | 0.23 | 0.33 | 0.14 | 0.00 | 0.00 | 0.00 |
| Patient 9 | 6.77 | IFN-γ | 0.45 | 0.15 | 0.12 | 0.51 | 0.24 | 0.11 | 1.49 | 0.00 | 0.97 |
| | | IL-2 | 0.16 | 0.21 | 0.16 | 0.16 | 0.24 | 0.16 | 0.00 | 0.00 | 0.00 |
| | | TNF-α | 0.24 | 0.15 | 0.14 | 0.34 | 0.20 | 0.24 | 2.99 | 0.00 | 0.00 |

| | | | IL-2/7/21 | | | | | | | | |
|---|---|---|---|---|---|---|---|---|---|---|---|
| | | | CD3 % | | | CD4 % | | | CD8 % | | |
| | Fold Expansion | | M | N | S | M | N | S | M | N | S |
| Patient 3 | 0.85 | IFN-γ | 0.97 | 0.35 | 0.23 | 1.27 | 0.44 | 0.18 | 0.00 | 0.00 | 0.00 |
| | | IL-2 | 0.73 | 0.66 | 0.31 | 1.27 | 1.10 | 0.37 | 0.00 | 0.00 | 0.52 |
| | | TNF-α | 1.70 | 0.31 | 0.17 | 2.55 | 0.88 | 0.18 | 0.00 | 0.72 | 0.00 |
| Patient 4 | 2.72 | IFN-γ | 0.21 | 0.38 | 0.29 | 0.07 | 0.16 | 0.09 | 0.00 | 0.33 | 0.72 |
| | | IL-2 | 0.09 | 0.05 | 0.09 | 0.17 | 0.00 | 0.09 | 0.40 | 0.33 | 0.00 |
| | | TNF-α | 0.13 | 0.19 | 0.16 | 0.17 | 0.22 | 0.18 | 0.40 | 0.00 | 0.00 |
| Patient 5 | 109.68 | IFN-γ | 0.87 | 0.06 | 0.47 | 1.62 | 0.09 | 1.11 | 1.52 | 0.00 | 0.54 |
| | | IL-2 | 0.07 | 0.07 | 0.07 | 0.20 | 0.12 | 0.14 | 0.00 | 0.00 | 0.00 |
| | | TNF-α | 0.18 | 0.07 | 0.20 | 0.44 | 0.20 | 0.59 | 0.76 | 0.57 | 0.54 |
| Patient 6 | 1.82 | IFN-γ | 0.21 | 0.17 | 0.13 | 0.18 | 0.00 | 0.00 | 0.28 | 0.21 | 0.34 |
| | | IL-2 | 0.16 | 0.05 | 0.11 | 0.45 | 0.51 | 0.00 | 0.10 | 0.07 | 0.34 |
| | | TNF-α | 0.19 | 0.22 | 0.14 | 0.45 | 1.01 | 2.58 | 0.94 | 0.35 | 0.93 |
| Patient 7 | 415.98 | IFN-γ | 0.74 | 0.30 | 0.50 | 2.93 | 0.81 | 1.36 | 0.39 | 0.39 | 0.41 |
| | | IL-2 | 0.10 | 0.07 | 0.15 | 0.43 | 0.22 | 0.14 | 0.39 | 0.13 | 0.00 |
| | | TNF-α | 0.44 | 0.18 | 0.25 | 2.22 | 0.81 | 0.61 | 0.39 | 0.26 | 0.10 |
| Patient 8 | 2.03 | IFN-γ | 0.14 | 0.21 | 0.11 | 0.23 | 0.09 | 0.11 | 0.00 | 5.82 | 0.00 |
| | | IL-2 | 0.06 | 0.05 | 0.11 | 0.10 | 0.12 | 0.20 | 1.75 | 0.53 | 0.83 |
| | | TNF-α | 0.13 | 0.25 | 0.26 | 0.25 | 0.33 | 0.44 | 0.00 | 0.53 | 0.42 |
| Patient 9 | 78.48 | IFN-γ | 0.76 | 0.25 | 0.49 | 1.87 | 0.30 | 1.01 | 0.00 | 0.60 | 1.11 |
| | | IL-2 | 0.03 | 0.11 | 0.09 | 0.13 | 0.56 | 0.57 | 0.00 | 0.60 | 0.37 |
| | | TNF-α | 0.19 | 0.14 | 0.25 | 0.52 | 0.19 | 0.64 | 0.00 | 0.60 | 0.00 |

Percent IFN-γ, IL-2 and TNF-α production from total CD3+, and in CD4+ and CD8+ subsets of COVID-19 reactive T cells derived from each recovered donor and stimulated with the peptide libraries derived from M, N and S structural proteins and fold expansion after culture with IL-2/4/21 or IL-2/7/21 cytokine cocktails.

### Table S4. Cytokine production of COVID-19 reactive T cells from recovered donors expanded with IL-2/4/7 or IL-2/7/15 against S1 and S2 (N and C terminals of the S protein), related to Figure 2.

| | | IL-2/4/7 | | | | | |
|---|---|---|---|---|---|---|---|
| | | CD3 % | | CD4 % | | CD8 % | |
| | | S1 | S2 | S1 | S2 | S1 | S2 |
| Patient 1 | IFN-γ | 4.69 | 0.71 | 4.47 | 0.66 | 0.70 | 0.60 |
| | IL-2 | 0.58 | 0.38 | 0.59 | 0.45 | 0.09 | 0.15 |
| | TNF-α | 7.19 | 1.11 | 5.97 | 0.75 | 0.26 | 0.45 |
| Patient 3 | IFN-γ | 1.33 | 5.12 | 1.00 | 0.46 | 1.43 | 3.62 |
| | IL-2 | 0.19 | 0.24 | 0.12 | 0.07 | 1.04 | 0.75 |
| | TNF-α | 2.62 | 0.92 | 1.86 | 0.51 | 7.04 | 1.09 |
| Patient 4 | IFN-γ | 10.20 | 7.84 | 8.33 | 6.60 | 13.20 | 6.29 |
| | IL-2 | 2.58 | 4.48 | 1.04 | 2.72 | 6.15 | 6.47 |
| | TNF-α | 2.85 | 1.52 | 2.30 | 0.88 | 3.71 | 1.15 |
| Patient 5 | IFN-γ | 26.20 | 1.93 | 9.10 | 2.14 | 12.20 | 3.12 |
| | IL-2 | 0.14 | 0.83 | 0.09 | 0.37 | 0.00 | 3.29 |
| | TNF-α | 4.62 | 2.63 | 3.62 | 1.60 | 9.38 | 2.44 |
| Patient 6 | IFN-γ | 6.92 | 1.09 | 10.60 | 1.31 | 2.70 | 1.05 |
| | IL-2 | 1.50 | 1.22 | 1.20 | 1.49 | 2.52 | 2.44 |
| | TNF-α | 9.47 | 0.72 | 12.80 | 0.86 | 0.00 | 0.00 |
| Patient 7 | IFN-γ | 26.50 | 8.54 | 20.80 | 3.66 | 12.90 | 2.11 |
| | IL-2 | 5.03 | 0.36 | 2.55 | 0.13 | 0.60 | 5.79 |
| | TNF-α | 2.42 | 1.30 | 1.67 | 0.61 | 8.70 | 2.63 |
| Patient 8 | IFN-γ | 1.53 | 3.84 | 1.42 | 1.34 | 1.88 | 3.40 |
| | IL-2 | 0.17 | 0.41 | 0.06 | 0.15 | 3.56 | 1.99 |
| | TNF-α | 3.62 | 1.67 | 2.81 | 1.08 | 27.30 | 13.70 |
| Patient 9 | IFN-γ | 2.67 | 3.92 | 1.22 | 1.76 | 3.34 | 3.60 |
| | IL-2 | 0.48 | 0.61 | 0.22 | 0.38 | 6.05 | 4.32 |
| | TNF-α | 0.45 | 1.63 | 0.33 | 1.00 | 3.97 | 14.70 |

| | | IL-2/7/15 | | | | | |
|---|---|---|---|---|---|---|---|
| | | CD3 % | | CD4 % | | CD8 % | |
| | | S1 | S2 | S1 | S2 | S1 | S2 |
| Patient 1 | IFN-γ | 5.96 | 3.25 | 4.47 | 3.72 | 0.83 | 7.77 |
| | IL-2 | 0.8 | 0.58 | 0.59 | 0.69 | 4.13 | 3.88 |
| | TNF-α | 7.59 | 1.17 | 5.97 | 0.93 | 5.79 | 0.97 |
| Patient 3 | IFN-γ | 7.43 | 1.61 | 1.05 | 0.76 | 4.57 | 1.72 |
| | IL-2 | 0.41 | 0.13 | 0.07 | 0.04 | 3.28 | 0.54 |
| | TNF-α | 5.06 | 2.48 | 3.70 | 1.45 | 26.20 | 18.40 |
| Patient 4 | IFN-γ | 18.00 | 14.60 | 2.50 | 1.32 | 10.10 | 5.90 |
| | IL-2 | 1.57 | 1.29 | 0.22 | 0.12 | 1.04 | 0.75 |
| | TNF-α | 1.94 | 1.93 | 1.53 | 1.31 | 8.33 | 8.86 |
| Patient 5 | IFN-γ | 4.99 | 1.96 | 0.82 | 1.81 | 5.86 | 0.46 |
| | IL-2 | 0.12 | 0.05 | 0.05 | 0.03 | 3.15 | 0.46 |
| | TNF-α | 1.63 | 4.14 | 0.84 | 3.05 | 12.20 | 22.60 |
| Patient 6 | IFN-γ | 3.95 | 2.72 | 1.97 | 2.53 | 1.22 | 1.20 |
| | IL-2 | 0.80 | 0.24 | 0.21 | 0.09 | 1.28 | 0.40 |
| | TNF-α | 2.46 | 2.29 | 1.51 | 1.41 | 18.70 | 15.30 |
| Patient 7 | IFN-γ | 11.10 | 1.72 | 15.80 | 2.00 | 2.70 | 1.05 |
| | IL-2 | 4.37 | 3.80 | 3.02 | 2.34 | 3.60 | 3.48 |
| | TNF-α | 10.50 | 0.95 | 13.90 | 1.03 | 0.36 | 0.00 |
| Patient 8 | IFN-γ | 1.73 | 4.34 | 2.71 | 10.50 | 1.81 | 3.70 |
| | IL-2 | 0.12 | 4.97 | 0.18 | 1.13 | 0.14 | 4.48 |
| | TNF-α | 1.57 | 2.10 | 1.38 | 2.25 | 16.10 | 13.90 |
| Patient 9 | IFN-γ | 5.92 | 5.03 | 7.23 | 6.43 | 4.48 | 2.71 |
| | IL-2 | 1.20 | 1.21 | 0.47 | 0.42 | 2.90 | 2.98 |
| | TNF-α | 1.59 | 1.41 | 4.48 | 0.38 | 0.40 | 0.27 |

Percent IFN-γ, IL-2 and TNF-α production from total CD3+, and CD4+ and CD8+ subsets of COVID-19 reactive T cells derived from each recovered donor and stimulated with the peptide libraries derived from S1 and S2 (N and C terminals of the S protein) after expansion with IL-2/4/7 or IL-2/7/15 cytokine cocktails.

**Table S5. Cytokine production of COVID-19 reactive T cells from recovered donors expanded with IL-2/4/21 or IL-2/7/21 against S1 and S2 (N and C terminals of the S protein), related to Figure 2.**

| | | IL-2/4/21 | | | | | |
|---|---|---|---|---|---|---|---|
| | | CD3 % | | CD4 % | | CD8 % | |
| | | S1 | S2 | S1 | S2 | S1 | S2 |
| Patient 3 | IFN-γ | 0.04 | 0.08 | 0.02 | 0.05 | 0.00 | 0.00 |
| | IL-2 | 0.12 | 0.16 | 0.13 | 0.16 | 0.00 | 0.00 |
| | TNF-α | 0.08 | 0.05 | 0.08 | 0.04 | 0.00 | 1.59 |
| Patient 4 | IFN-γ | 0.17 | 0.19 | 0.14 | 0.23 | 0.00 | 0.00 |
| | IL-2 | 0.40 | 0.57 | 0.68 | 0.23 | 0.00 | 0.00 |
| | TNF-α | 0.40 | 0.19 | 0.27 | 0.00 | 0.00 | 0.00 |
| Patient 5 | IFN-γ | 4.15 | 0.14 | 6.21 | 0.22 | 0.00 | 0.00 |
| | IL-2 | 0.17 | 0.08 | 0.24 | 0.09 | 0.00 | 0.00 |
| | TNF-α | 1.78 | 0.13 | 2.69 | 0.22 | 0.00 | 0.00 |
| Patient 6 | IFN-γ | 0.33 | 0.14 | 0.17 | 0.22 | 0.00 | 0.00 |
| | IL-2 | 0.00 | 0.57 | 0.00 | 0.44 | 0.00 | 0.00 |
| | TNF-α | 1.00 | 0.72 | 0.34 | 0.22 | 0.00 | 0.00 |
| Patient 7 | IFN-γ | 0.68 | 0.21 | 1.12 | 0.60 | 0.67 | 0.00 |
| | IL-2 | 0.21 | 0.18 | 0.31 | 0.22 | 0.22 | 0.33 |
| | TNF-α | 0.23 | 0.12 | 0.35 | 0.18 | 0.00 | 0.67 |
| Patient 8 | IFN-γ | 0.08 | 0.09 | 0.00 | 0.14 | 0.00 | 0.00 |
| | IL-2 | 0.35 | 0.16 | 0.14 | 0.00 | 0.66 | 0.61 |
| | TNF-α | 0.18 | 0.14 | 0.29 | 0.00 | 0.00 | 0.00 |
| Patient 9 | IFN-γ | 0.03 | 0.26 | 0.08 | 0.50 | 0.00 | 0.00 |
| | IL-2 | 0.10 | 0.13 | 0.11 | 0.12 | 0.00 | 0.00 |
| | TNF-α | 0.07 | 0.20 | 0.16 | 0.30 | 0.00 | 0.00 |
| | | IL-2/7/21 | | | | | |
| | | CD3 % | | CD4 % | | CD8 % | |
| | | S1 | S2 | S1 | S2 | S1 | S2 |
| Patient 3 | IFN-γ | 0.05 | 0.10 | 0.00 | 0.00 | 0.58 | 0.00 |
| | IL-2 | 0.11 | 0.21 | 0.09 | 0.43 | 0.00 | 0.69 |
| | TNF-α | 0.22 | 0.07 | 0.35 | 0.14 | 0.00 | 0.00 |
| Patient 4 | IFN-γ | 0.20 | 0.13 | 0.23 | 0.10 | 0.00 | 0.00 |
| | IL-2 | 0.07 | 0.21 | 0.10 | 0.24 | 0.00 | 0.87 |
| | TNF-α | 0.16 | 0.15 | 0.20 | 0.07 | 0.00 | 0.00 |
| Patient 5 | IFN-γ | 0.89 | 0.05 | 2.22 | 0.08 | 0.00 | 0.00 |
| | IL-2 | 0.08 | 0.14 | 0.17 | 0.11 | 0.00 | 0.83 |
| | TNF-α | 0.37 | 0.06 | 1.01 | 0.15 | 0.00 | 0.83 |
| Patient 6 | IFN-γ | 0.20 | 0.13 | 0.34 | 1.00 | 0.43 | 0.10 |
| | IL-2 | 0.06 | 0.08 | 0.34 | 1.50 | 0.22 | 0.00 |
| | TNF-α | 0.16 | 0.07 | 1.02 | 1.50 | 0.43 | 0.50 |
| Patient 7 | IFN-γ | 0.31 | 0.12 | 0.86 | 0.21 | 0.63 | 0.47 |
| | IL-2 | 0.06 | 0.04 | 0.11 | 0.39 | 0.00 | 0.00 |
| | TNF-α | 0.20 | 0.09 | 1.18 | 0.21 | 0.63 | 0.00 |
| Patient 8 | IFN-γ | 0.10 | 0.09 | 0.13 | 0.05 | 0.56 | 0.00 |
| | IL-2 | 0.08 | 0.07 | 0.18 | 0.18 | 0.00 | 0.00 |
| | TNF-α | 0.20 | 0.23 | 0.31 | 0.55 | 0.00 | 0.00 |
| Patient 9 | IFN-γ | 0.05 | 0.63 | 0.04 | 1.24 | 0.27 | 0.35 |
| | IL-2 | 0.08 | 0.07 | 0.37 | 0.41 | 0.54 | 0.35 |
| | TNF-α | 0.08 | 0.31 | 0.20 | 0.85 | 0.54 | 0.35 |

Percent IFN-γ, IL-2 and TNF-α production from total CD3+, and CD4+ and CD8+ subsets of COVID-19 reactive T cells derived from each recovered donor and stimulated with the peptide libraries derived from S1 and S2 (N and C terminals of the S protein) after expansion with IL-2/4/21 or IL-2/7/21 cytokine cocktails.

**Table S6. Cytokine production of COVID-19 reactive T cells from healthy controls expanded with different cytokine cocktails against M, N and S structural proteins, related to Figure 4.**

| | | CD3 % | | | CD4 % | | | CD8 % | | |
|---|---|---|---|---|---|---|---|---|---|---|
| | | M | N | S | M | N | S | M | N | S |
| IL-2/4/7 | IFN-γ median | 0.43 | 0.29 | 0.44 | 0.55 | 0.28 | 0.34 | 0.11 | 0.18 | 0.24 |
| | IFN-γ min | 0.13 | 0.16 | 0.19 | 0.14 | 0.16 | 0.06 | 0.02 | 0.07 | 0.04 |
| | IFN-γ max | 1.37 | 0.67 | 2.57 | 2.59 | 1.12 | 2.50 | 0.19 | 0.67 | 2.59 |
| | IL-2 median | 0.14 | 0.50 | 0.16 | 0.10 | 0.16 | 0.10 | 0.24 | 0.18 | 0.24 |
| | IL-2 min | 0.03 | 0.12 | 0.04 | 0.03 | 0.14 | 0.04 | 0.00 | 0.09 | 0.02 |
| | IL-2 max | 0.31 | 0.74 | 0.63 | 0.23 | 1.27 | 0.17 | 0.41 | 0.63 | 0.29 |
| | TNF-α median | 0.25 | 0.23 | 0.16 | 0.26 | 0.57 | 0.27 | 0.04 | 0.22 | 0.19 |
| | TNF-α min | 0.15 | 0.17 | 0.06 | 0.11 | 0.07 | 0.07 | 0.02 | 0.12 | 0.02 |
| | TNF-α max | 1.11 | 2.34 | 1.59 | 2.19 | 5.17 | 3.55 | 0.18 | 0.28 | 0.67 |
| IL-2/7/15 | IFN-γ median | 0.24 | 0.15 | 0.59 | 0.27 | 0.18 | 0.37 | 0.10 | 0.13 | 0.11 |
| | IFN-γ min | 0.10 | 0.09 | 0.21 | 0.17 | 0.05 | 0.16 | 0.07 | 0.08 | 0.06 |
| | IFN-γ max | 2.13 | 0.39 | 1.64 | 6.63 | 0.49 | 1.58 | 0.91 | 0.24 | 1.22 |
| | IL-2 median | 0.17 | 0.12 | 0.18 | 0.10 | 0.12 | 0.11 | 0.11 | 0.08 | 0.15 |
| | IL-2 min | 0.05 | 0.00 | 0.07 | 0.09 | 0.00 | 0.07 | 0.02 | 0.00 | 0.07 |
| | IL-2 max | 0.25 | 0.25 | 0.29 | 0.25 | 0.14 | 0.31 | 0.22 | 0.21 | 0.27 |
| | TNF-α median | 0.12 | 0.12 | 0.18 | 0.24 | 0.21 | 0.31 | 0.10 | 0.10 | 0.14 |
| | TNF-α min | 0.09 | 0.02 | 0.09 | 0.10 | 0.00 | 0.14 | 0.06 | 0.01 | 0.08 |
| | TNF-α max | 1.44 | 0.14 | 0.38 | 4.88 | 0.23 | 1.14 | 0.24 | 0.18 | 0.22 |

Percent IFN-γ, IL-2 and TNF-α production (median, minimum and maximum values) from total CD3+, and CD4+ and CD8+ subsets of COVID-19 reactive T cells stimulated with the peptide libraries derived from M, N and S structural proteins after expansion with the two different cytokine cocktails IL-2/4/7 or IL-2/7/15.

**Table S7. Cytokine production and fold expansion of COVID-19 reactive T cells derived from the different healthy controls and expanded with IL-2/4/7 or IL-2/7/15 cytokine cocktails, related to Figure 4.**

| | | | IL-2/4/7 | | | | | | | | |
|---|---|---|---|---|---|---|---|---|---|---|---|
| | | | CD3 % | | | CD4 % | | | CD8 % | | |
| | Fold Expansion | | M | N | S | M | N | S | M | N | S |
| HD 1 | 21.31 | IFN-γ | 0.14 | 0.29 | 2.57 | 0.17 | 0.32 | 2.16 | 0.02 | 0.07 | 0.04 |
| | | IL-2 | 0.31 | 0.74 | 0.63 | 0.11 | 0.14 | 0.07 | 0.26 | 0.63 | 0.29 |
| | | TNF-α | 0.15 | 0.23 | 0.06 | 0.14 | 0.27 | 0.58 | 0.02 | 0.22 | 0.11 |
| HD 2 | 8.13 | IFN-γ | 0.13 | 0.25 | 0.44 | 0.14 | 0.28 | 0.24 | 0.19 | 0.30 | 0.22 |
| | | IL-2 | 0.25 | 0.50 | 0.27 | 0.10 | 0.16 | 0.10 | 0.41 | 0.30 | 0.27 |
| | | TNF-α | 0.16 | 0.21 | 0.16 | 0.26 | 0.57 | 0.27 | 0.18 | 0.27 | 0.19 |
| HD 3 | 2.85 | IFN-γ | 0.43 | 0.16 | 0.19 | 0.55 | 0.16 | 0.06 | 0.11 | 0.18 | 0.24 |
| | | IL-2 | 0.14 | 0.12 | 0.13 | 0.23 | 0.15 | 0.10 | 0.24 | 0.18 | 0.06 |
| | | TNF-α | 0.25 | 0.17 | 0.12 | 0.11 | 0.07 | 0.07 | 0.17 | 0.12 | 0.38 |
| HD4 | 27.74 | IFN-γ | 1.37 | 0.67 | 0.33 | 1.57 | 1.12 | 0.34 | 0.08 | 0.17 | 0.41 |
| | | IL-2 | 0.04 | 0.65 | 0.04 | 0.04 | 1.27 | 0.04 | 0.00 | 0.15 | 0.02 |
| | | TNF-α | 1.11 | 2.34 | 0.22 | 1.38 | 5.17 | 0.27 | 0.03 | 0.28 | 0.02 |
| HD5 | 41.84 | IFN-γ | 0.84 | 0.60 | 1.83 | 2.59 | 0.27 | 2.50 | 0.12 | 0.67 | 2.59 |
| | | IL-2 | 0.03 | 0.15 | 0.16 | 0.03 | 0.22 | 0.17 | 0.03 | 0.09 | 0.24 |
| | | TNF-α | 0.63 | 0.62 | 1.59 | 2.19 | 0.96 | 3.55 | 0.04 | 0.22 | 0.67 |

| | | | IL-2/7/15 | | | | | | | | |
|---|---|---|---|---|---|---|---|---|---|---|---|
| | | | CD3 % | | | CD4 % | | | CD8 % | | |
| | Fold Expansion | | M | N | S | M | N | S | M | N | S |
| HD 1 | 15.73 | IFN-γ | 0.24 | 0.23 | 0.56 | 0.26 | 0.30 | 0.36 | 0.10 | 0.10 | 0.11 |
| | | IL-2 | 0.25 | 0.25 | 0.29 | 0.09 | 0.12 | 0.09 | 0.22 | 0.18 | 0.18 |
| | | TNF-α | 0.09 | 0.12 | 0.09 | 0.13 | 0.21 | 0.23 | 0.07 | 0.10 | 0.08 |
| HD 2 | 9.01 | IFN-γ | 0.10 | 0.11 | 0.21 | 0.17 | 0.13 | 0.16 | 0.07 | 0.17 | 0.06 |
| | | IL-2 | 0.17 | 0.13 | 0.18 | 0.10 | 0.12 | 0.11 | 0.15 | 0.21 | 0.27 |
| | | TNF-α | 0.12 | 0.14 | 0.15 | 0.24 | 0.22 | 0.34 | 0.10 | 0.13 | 0.11 |
| HD 3 | 4.00 | IFN-γ | 0.24 | 0.15 | 0.59 | 0.27 | 0.18 | 0.37 | 0.10 | 0.08 | 0.09 |
| | | IL-2 | 0.13 | 0.12 | 0.14 | 0.13 | 0.14 | 0.18 | 0.10 | 0.08 | 0.13 |
| | | TNF-α | 0.12 | 0.13 | 0.18 | 0.10 | 0.11 | 0.14 | 0.22 | 0.18 | 0.15 |
| HD4 | 53.95 | IFN-γ | 2.13 | 0.09 | 0.67 | 6.63 | 0.05 | 0.40 | 0.21 | 0.24 | 0.96 |
| | | IL-2 | 0.05 | 0.00 | 0.07 | 0.10 | 0.00 | 0.07 | 0.02 | 0.00 | 0.07 |
| | | TNF-α | 1.44 | 0.02 | 0.33 | 4.88 | 0.00 | 0.31 | 0.06 | 0.07 | 0.22 |
| HD5 | 24.77 | IFN-γ | 1.23 | 0.39 | 1.64 | 1.24 | 0.49 | 1.58 | 0.91 | 0.13 | 1.22 |
| | | IL-2 | 0.17 | 0.00 | 0.18 | 0.25 | 0.01 | 0.31 | 0.11 | 0.01 | 0.15 |
| | | TNF-α | 0.47 | 0.07 | 0.38 | 1.06 | 0.23 | 1.14 | 0.24 | 0.01 | 0.14 |

Percent IFN-γ, IL-2 and TNF-α production from total CD3+, and CD4+ and CD8+ subsets of COVID-19 reactive T cells derived from each healthy donor and stimulated with the peptide libraries derived from M, N and S structural proteins and fold expansion after culture with IL-2/4/7 or IL-2/7/15 cytokine cocktails.
